# Modelling Speed-Accuracy Tradeoffs in the Stopping Rule for Confidence Judgments

**DOI:** 10.1101/2023.02.27.530208

**Authors:** Stef Herregods, Pierre Le Denmat, Luc Vermeylen, Kobe Desender

## Abstract

Making a decision and reporting confidence in the accuracy of that decision are thought to be driven by the same mechanism: the accumulation of evidence. Previous research has shown that choices and reaction times are well accounted for by a computational model assuming noisy accumulation of evidence until crossing a decision boundary (e.g., the drift diffusion model). Decision confidence can be derived from the amount of evidence following post-decision evidence accumulation. Currently, the stopping rule for post-decision evidence accumulation is underspecified. In the current work, we quantitatively and qualitatively compare the ability of four prominent models of confidence couched within evidence accumulation to account for this stopping rule. In two experiments, participants were instructed to make fast or accurate decisions, and to give fast or carefully considered confidence judgments. We then compared the different models in their ability to capture the speed-accuracy effects on confidence. Both qualitatively and quantitatively, the data were best accounted for by our newly proposed Flexible Collapsing Boundaries model, in which post-decision accumulation terminates once it reaches one of two opposing slowly collapsing confidence boundaries. Inspection of the parameters of this model revealed that instructing participants to make fast versus accurate decisions influenced the height of the decision boundaries, while instructing participants to make fast versus careful confidence judgments influenced height of the confidence boundaries. Our data show that the stopping rule for confidence judgments can be well described as an accumulation-to-bound process, and that the height of these confidence boundaries are under strategic control.

## Introduction

Human decision making is accompanied by a sense of confidence. Humans often report high confidence when they make correct decisions and low confidence when they make incorrect decisions (Fleming et al., 2010). Understanding the computational underpinnings of decision confidence is of high importance, given that humans use decision confidence to adapt subsequent behavior (Desender et al., 2018, 2019; Folke et al., 2016). In recent work, identifying the computational underpinnings of decision confidence has been identified as an important common goal for the field of metacognition (Rahnev et al., 2022). Given that decision confidence reflects an evaluation of the accuracy of a decision, computational accounts of decision confidence usually depart from decision making models and aim to explain the computation of confidence within these models.

In many decision-making scenarios, human observers face the challenging task to make accurate decisions based on noisy evidence. Many theories of decision making assume that people solve this challenge by accumulating multiple pieces of evidence. Accumulation-to-bound models specifically propose that evidence is accumulated sequentially until the accumulated evidence reaches a predefined decision boundary. Once the decision boundary is reached, the model makes a choice (for review, see Gold & Shadlen, 2007; Figure 1A). Within the drift diffusion model (DDM), evidence accumulates towards one of two opposing decision boundaries, with the additional assumption that evidence for both choice options is perfectly anti-correlated (Ratcliff & McKoon, 2008). In its most basic implementation, the DDM explains the dynamics of decision making using only three main parameters: a drift rate, reflecting the strength of the evidence accumulation process, a decision boundary, reflecting the degree of evidence required before a decision is made, and non-decision time, capturing non-decision related components. This simple tenet has proven to be a powerful framework that can account for a realm of behavioral and neurophysiological data. For example, accumulation-to-bound signals such as described by the DDM have been observed in human (Donner et al., 2009; O’Connell et al., 2012) and primate (Gold & Shadlen, 2007) neurophysiology. Most prominently, the DDM can explain the tradeoff between speed and accuracy that characterizes all forms of speeded decision making (Bogacz et al., 2006; Bogacz, Wagenmakers, et al., 2010). When participants are instructed to make speeded versus accurate decisions, the DDM explains these data by changing the height of the decision boundary (although the selectivity of this effect has been debated; Rafiei & Rahnev, 2021). Decreasing the decision boundary effectively lowers the required level of evidence before reaching it, promoting fast responses at the expense of accuracy. Given that participants are able to change the decision boundary based on instructions (amongst many other manipulations), it is believed that the height of the decision boundary is under voluntary strategic control (Balci et al., 2011; Bogacz, Hu, et al., 2010).

**Figure 1.**
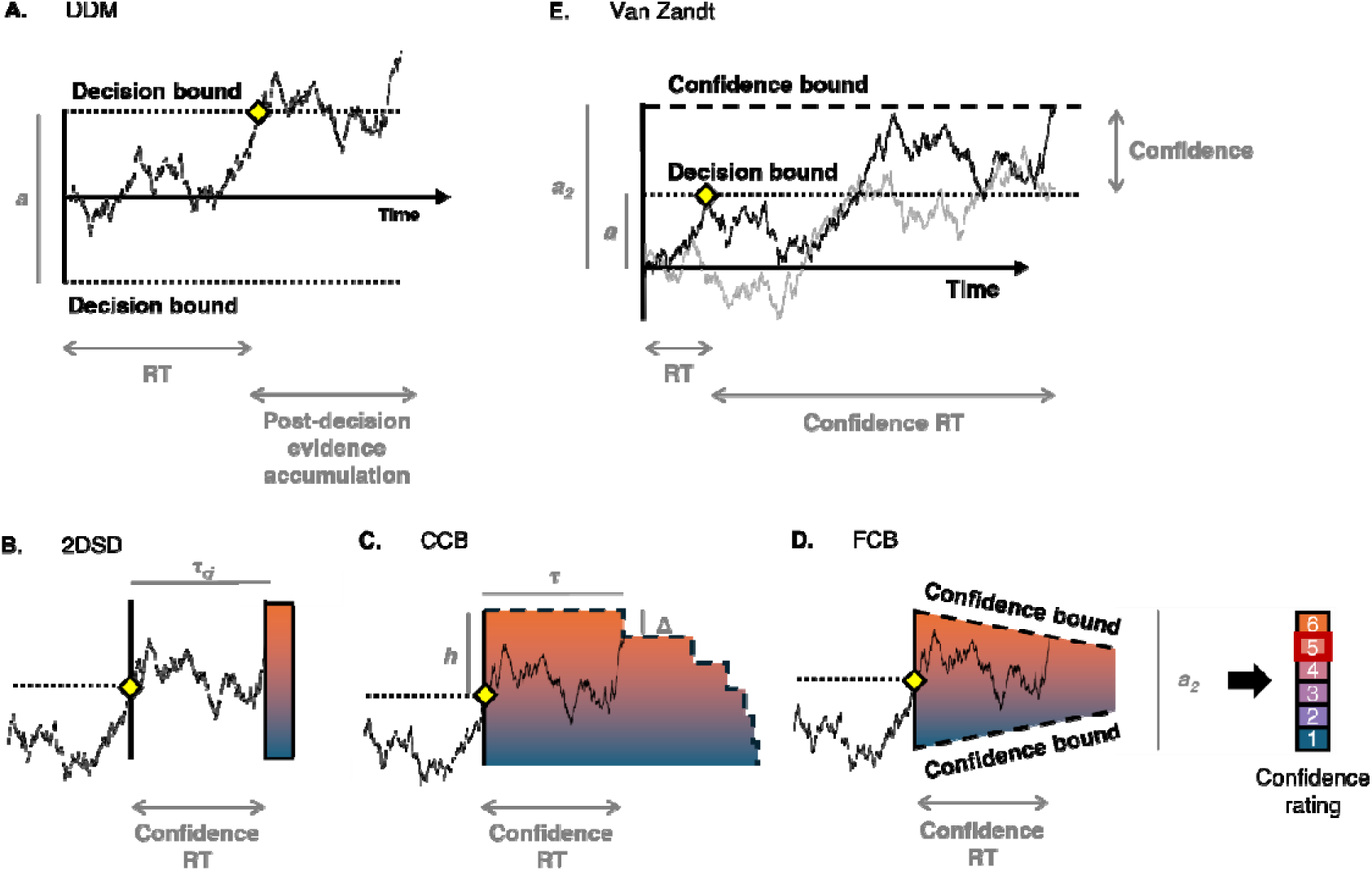
Competing models implementing post-decision confidence computation. (A) According to the DDM, evidence accumulates until reaching a decision boundary at which point a decision is made, here indicated by the yellow diamond. Boundary separation (a) can vary, changing the speed-accuracy trade-off. Confidence is often modelled by allowing the accumulation process to continue after boundary crossing, however the exact nature of the stopping rule for post-decision processing is unclear. (B) The 2DSD model proposes that post-decision evidence accumulates for a fixed time period (τ) after making the decision. The final state of the post-decision accumulator then informs the confidence rating. (C) The CCB model assumes that the accumulation of post-decision evidence terminates once it reaches a slowly collapsing confidence boundary, with absolute height (h), collapse time (τ) and collapse height (Δ) as free parameters. (D) In the newly proposed FCB model, post-decision evidence accumulates until reaching one of two slowly collapsing confidence boundaries, with confidence boundary separation a_2_ and confidence urgency u_2_. (E) The race model proposed by Van Zandt and Maldonado-Molina (2004) assumes separate evidence accumulators for both decision options. Confidence is quantified as the difference in evidence once one of both accumulators reaches the confidence boundary, with boundary height a2.

Given the success of the DDM in explaining decision making, several attempts have been made to explain decision confidence within this model. Capitalizing on the notion that the sense of confidence seems to arise *after* a decision has been made, Pleskac and Busemeyer (2010) put forward the two-stage dynamic signal detection theory model (2DSD) which proposes that the process of evidence accumulation does not terminate once a choice boundary has been crossed, but rather there is continued accumulation of (post-decision) evidence, which determines decision confidence. If additional post-decision evidence confirms the initial decision, the model will produce a high confidence response. If additional post-decision evidence contradicts the initial decision, the model produces low confidence, or even changes its mind about the initial decision (Resulaj et al., 2009; Van Den Berg et al., 2016). Given that post-decision evidence is most likely to contradict initial decisions when these were incorrect, this account can explain why confidence is usually higher for correct than for incorrect decisions (Moran et al., 2015; Pleskac & Busemeyer, 2010), and why confidence better tracks accuracy when participants take more time to report confidence (Yu et al., 2015).

Previous modeling work has shown that the 2DSD model can jointly explain choices, reaction times and decision confidence (see also Calder-Travis et al., 2024; Hellmann et al., 2023; van den Berg et al., 2016; Zylberberg et al., 2016). Strikingly, much less attention has been devoted towards the speed with which confidence reports are provided. This is remarkable, given that confidence RTs are highly informative about the underlying computations (Moran et al., 2015). As a consequence, models such as the 2DSD propose a very simplistic stopping rule for the post-decision evidence accumulation process (see also Desender et al., 2022; Hellmann et al., 2023; Yu et al., 2015) or do not contain an explicit stopping rule at all (Balsdon et al., 2020; Pereira et al., 2021, 2022). Within the 2DSD model, an additional parameter is included which controls the duration of the post-decision processing time (i.e. the time between the choice and the confidence report). Thus, in the 2DSD the stopping rule for confidence judgments is to stop accumulating post-decision evidence once a certain amount of time has passed (Figure 1B). At first sight, such a static implementation seems incompatible with the considerable variation in confidence RTs that is usually observed in empirical data. Indeed, under a strict interpretation, this account predicts that confidence judgments will always be provided after a fixed latency, potentially with some variability. Contrary to this, confidence RTs show the same right-skewed distributions as choice RTs. Importantly, although it thus seems that the 2DSD provides an unrealistic description of the stopping rule for post-decision accumulation, to our knowledge a comprehensive model fitting exercise evaluating this critique and quantitative comparison with competing models to unravel more appropriate mechanisms has not yet been performed. Therefore, in the current work we set out to do exactly this and compare the ability of 2DSD to several other candidate models in explaining the stopping rule for confidence judgments.

Apart from the 2DSD, a first competing model that will be considered is the Collapsing Confidence Boundary model (CCB) put forward by Moran and colleagues (2015). In line with the observation that confidence RTs provide reliable information about the stopping rule of confidence, Moran and colleagues (2015) proposed a model with continuous post-decision accumulation until reaching a single slowly collapsing confidence boundary (Figure 1C). The authors showed that the CCB model was able to account for a variety of empirical patterns involving confidence RTs and confidence judgments, which could not be accounted for by the 2DSD. The work from Moran et al. (2015) suggests a (collapsing) confidence boundary as the stopping rule for confidence judgments.

A second competing model that will be considered has been proposed by Van Zandt and Maldonado-Molina, 2004 in an effort to explain response reversals in recognition memory. In short, these authors propose a race between two fully independent evidence accumulators towards a choice boundary (determining the choice and a corresponding RT), and subsequently towards a confidence boundary (determining the confidence RT). The level of confidence is then quantified as the distance between both accumulators at the moment of reaching the confidence boundary (Figure 1E). This model is similar in spirit to the CCB put forward by Moran et al. (2015) except that it posits the existence of a flat confidence boundary (as opposed to a collapsing confidence boundary), that it quantifies confidence as the balance-of-evidence between both accumulators (see also Vickers, 1979), and that it proposes a race between two fully independent accumulators (as opposed to having full inhibition between both accumulators, as in the drift diffusion model).

Finally, we propose a third competing model which extends the CCB approach such that it allows for more flexibility in capturing various association between confidence and confidence RTs. Specifically, due to its architecture the CCB always predicts that longer confidence RTs are associated with lower confidence (see Figure 1C). Such a pattern is quite common in experiments where participants report their confidence on scale running from uncertain until certain. However, as shown below, in experiments where participants are asked to report their confidence on a scale that also includes the possibility to detect their own errors (i.e., running from certainly wrong until certainly correct), sometimes the relation between confidence RTs and confidence shows an inverted U-shape. To account for such pattern, we here propose a model in which both choice and confidence are implemented as a process of continuous accumulation until one of two opposing boundaries is reached. For convenience, we refer to this model as the Flexible Confidence Boundary (FCB) model (Figure 1D).

In the current work, we compared the ability of these four models to accurately describe the stopping rule for confidence judgments, by fitting them to data from two experiments (one with a binary and one with a 6-choice confidence report). To provide a strong experimental manipulation of the stopping rule, in both experiments we manipulated the level of caution that participants used to provide their confidence reports. Specifically, just like how individuals can modulate their choice boundaries according to speed-accuracy tradeoff instructions (Ratcliff & McKoon, 2008), we similarly instructed participants to either provide very fast confidence judgments or think carefully about their level of confidence. Remarkably, although there are numerous studies that have investigated speed-accuracy tradeoffs in choice formation (for review, see Bogacz, Wagenmakers, et al., 2010), to our knowledge it has yet to be investigated whether similar tradeoffs can be observed in confidence formation, and if so which of the above mentioned computational models of confidence can best account for such tradeoffs.

## Experiment 1 Methods and Materials

### Preregistration and Code

All hypotheses, sample sizes, exclusion criteria for participants, analyzed variables, the experimental design and planned analyses were preregistered on the Open Science Framework (OSF) registries (https://doi.org/10.17605/OSF.IO/Z2UCM), unless specified as exploratory. Additionally, all code and data are made publicly available on GitHub (https://github.com/StefHerregods/ConfidenceBounds).

### Participants

We decided *a priori* to test a minimum of 40 viable participants, in line with previous speed-accuracy trade-off research (Desender et al., 2022). Participant recruitment continued until this sample size was met after applying exclusion criteria. In total, 51 participants took part in Experiment 1 in return for course credit. From the total dataset, one participant gave the same confidence rating in more than 95% of the trials and 10 participants required too many training trials or did not complete the experiment in time. Data from these participants was excluded from further analyses. The final dataset comprised 40 participants (36 female), with a mean age of 18.0 (*SD* = 0.6, range = 17-19). All participants had normal or corrected-to-normal vision, and signed informed consent before their participation. The experiment was approved by the local ethics committee.

### Stimuli and Apparatus

The experiment was programmed using Python v3.6.6 and PsychoPy (Peirce et al., 2019). Participants completed the experiment on 24-inch LCD screens using an AZERTY-keyboard, with blue stickers indicating buttons used for confidence judgments and red stickers indicating decision-making buttons.

### Procedure

Each experimental trial started with the display of a white fixation cross on a black background for 1s (see Figure 2). Instructions regarding the speed-accuracy regime were shown above and below the fixation cross for decision-making and confidence judgments, respectively. Depending on the block, the instructions were to either ‘Make fast decisions’ or ‘Make accurate decisions’, and to ‘Give fast confidence ratings’ or ‘Think carefully about your confidence ratings’, for choices and confidence reports, respectively. For convenience, we will refer to both type of instructions as choice SAT and confidence SAT, respectively. Next, a dynamic random dot motion stimulus was presented until participants gave a response. If participants did not provide a response within 5s, the message “Too slow, please respond faster” was shown on the screen. Motion coherence was controlled by the proportion of dots consistently moving towards the left versus right side of the screen. During the main experiment, three levels of coherence were used (.1, .2 and .4). Participants were instructed to press the ‘c’ or ‘n’ key with the thumbs of their left and right hand, to indicate whether they thought dots were moving towards the left or the right, respectively. If participants responded within 5s, they were subsequently asked about their level of confidence. The text ‘How confident are you that you made the correct choice?’ appeared on top of the screen, and participants pressed the ‘e’ or the ‘u’ key with their index fingers, mapped to high and low confidence, respectively (mapping counterbalanced across participants). Confidence judgments were transformed to numeric values, with ‘low confidence’ as zero and ‘high confidence’ as one.

**Figure 2.**
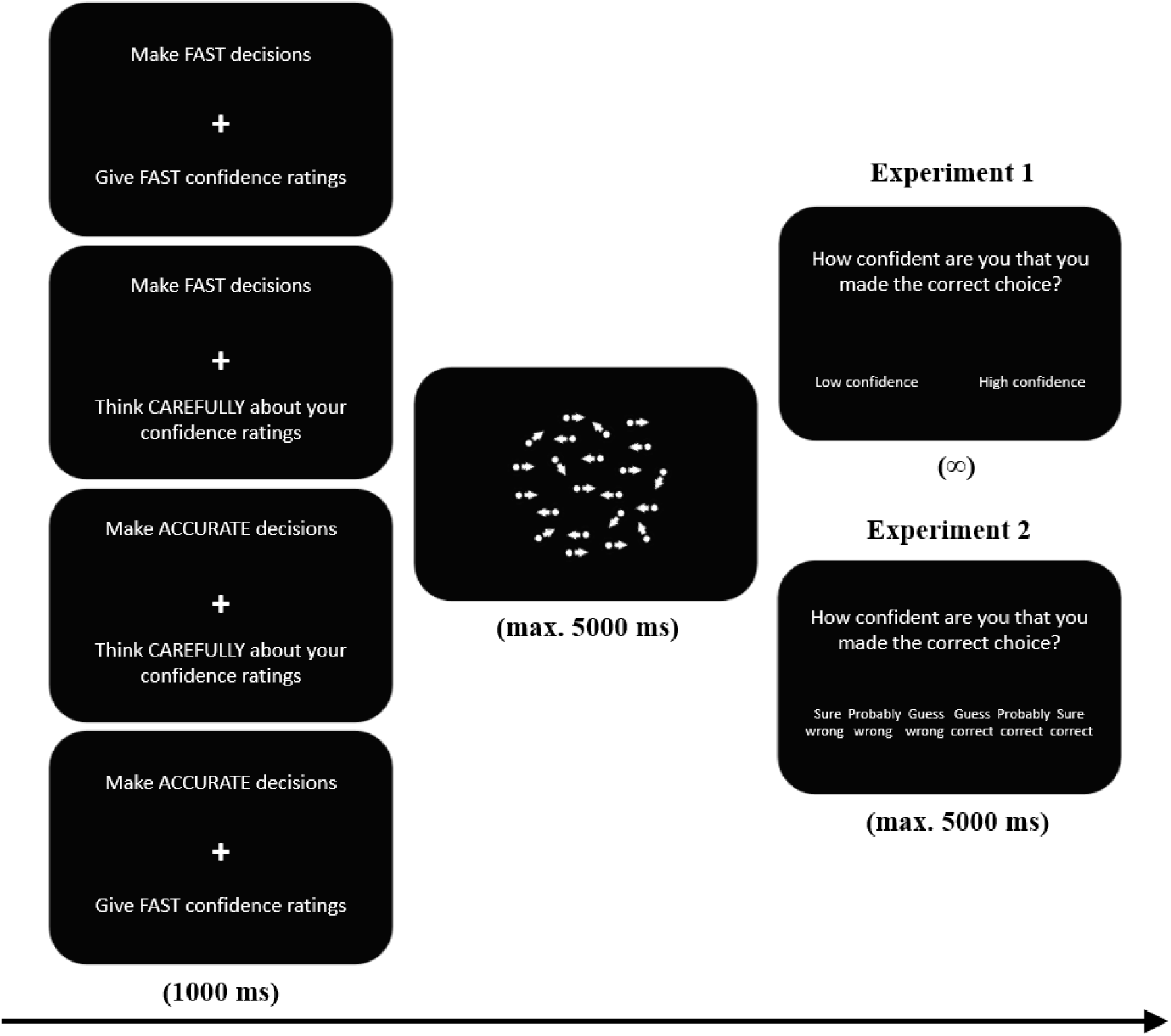
Example of an experimental trial. During presentation of the fixation cross participants received specific instructions regarding the speed-accuracy regime for choices (above fixation) and confidence (below fixation). These instructions were constant within a block, but switched each block. Next, participants made binary choices about random dot motion, and afterwards indicated their level of confidence on a two-point scale (Experiment 1), or a six-point scale (Experiment 2).

The experiment started with three practice blocks (24 trials each). In block 1 participants only made random dot motion decisions with a coherence of .5 for all trials. During this block they received feedback about choice accuracy after each trial. Participants repeated block 1 until achieving average accuracy of 85% or more. Block 2 was identical except that the same three coherence levels as in the main phase were used (.1, .2 and .4). Participants repeated block 2 until achieving average accuracy of 60% or more. In block 3, participants no longer received trial-by-trial feedback but instead were asked about their level of confidence after each trial. Afterwards, participants took part in twelve blocks of 60 trials each. In each block there was an equal number of coherent left and right dot motion trials, and an equal occurrence of the three coherence levels. Finally, each block had specific instructions about the speed-accuracy regime for decision-making and confidence judgments. These instructions appeared both before each block and at the start of each trial (i.e. during the fixation cross). Speed-accuracy regime instructions were constant within a block, but switched after each block. Each combination of instructions appeared three times, and the order of appearance was counterbalanced across participants using a Latin square. After each block, participants received feedback about their average accuracy, average reaction time and average confidence reaction time of the preceding block.

### Statistical analyses

RT’s and confidence RT’s on correct trials, accuracy, and confidence judgments on correct trials were analyzed using mixed effects models. All models included at least a random intercept per participant, and all manipulations (choice SAT, confidence SAT and coherence) and their interactions as fixed effects, unless otherwise specified. These models were then extended with random slopes in order of biggest increase in BIC, until the addition of random slopes led to a non-significant increase in likelihood or until the random effects structure was too complex to be supported by the data (leading to an unstable fit). We used the lmer and glmer functions of the lme4 package (Bates et al., 2015) to fit the linear and generalized linear mixed models, respectively, in R (R Core Team, 2021). The calculation of *p* values is based on chi-square estimations using the Wald test from the car-package (Fox & Weinberg, 2019). Due to violations of the assumptions of normally distributed residuals and homoscedasticity, all RT’s and confidence RT’s were log transformed and mean-centered. Furthermore, evidence accumulation model performance was compared across three different model specifications using BIC values computed as explained in Solway and Botvinick (2015). Finally, the influence of the speed-accuracy manipulations on the estimated model parameters of the best performing model was examined using a repeated measures ANOVA’s and follow-up paired t-tests, as implemented in the rstatix package (Kassambara, 2021).

### Model Specification

For the 2DSD, CCB and FCB models, we simulated noisy evidence accumulation using a random walk approximation of the drift diffusion process (Tuerlinckx et al., 2001). A random walk process started at *z*a*, with z being an unbiased starting point of .5, and continued to accumulate until the accumulated evidence reaches 0 or *a* (reflecting the height of the decision boundaries). At each time step τ the accumulated evidence was updated with Δ, with the update rule shown in equation (1):

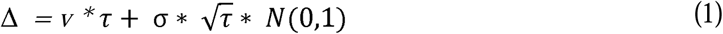

with *v* reflecting the drift rate, *N* reflecting the standard normal distribution, *ꚍ* reflecting precision, which was set to .001 in all simulations, and σ reflecting within-trial noise which was fixed to 1. Choice and RT were quantified at the moment of boundary crossing. We used an accuracy coding scheme, such that crossing the upper decision boundary equals a correct choice and crossing the lower decision boundary an incorrect choice. An additional time *ter* was added to predicted RTs to capture non-decision-related processes. After the accumulated evidence reached 0 or *a*, evidence continued to accumulate at each time step *ꚍ* with displacement *Δp*, with the post-decision update rule shown in equation (2):

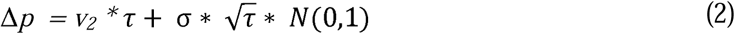

with v_2_ reflecting the drift rate governing post-decisional processing. Allowing dissociations between drift rate and post-decisional drift rate is necessary to account for differences in metacognitive accuracy (Desender et al., 2022). Post-decisional accumulation continued until a certain criterion was reached, and this stopping rule differed across the models we investigated.

For the first model, based on the 2DSD model of Pleskac and Busemeyer (2010), evidence continues to accumulate post-decision for a variable inter-judgment time, normally distributed with mean τ_cj_ and standard deviation SD_τ_. Confidence judgments are then determined by the crossing of confidence criterion *c* + *a* for correct decision trials or -*c* for incorrect decision trials. More specifically, higher evidence than the confidence criterion towards the chosen decision leads to a ‘high confidence’ response, and less evidence to a ‘low confidence’ response. Note that in the original 2DSD model proposed by Pleskac and Busemeyer (2010), across-trial variability in drift rate and starting point variability were also modelled, but we opted to not implement these parameters such that all the different models considered here are similar in their decision process and only differed in the post-decision stopping mechanisms.

In the second model, based on the CCB model of Moran et al. (2015), evidence accumulates until it reaches a single “stepwise” collapsing confidence boundary (i.e. an evidence-based criterion) with the starting height being determined by parameter *h.* When the accumulator reaches the confidence boundary at height *h* the model gives a high confidence response. After collapse time τ_cj_, the criterion height is lowered with collapse height Δ. In the variant with multiple discrete confidence levels, predicted model confidence lowers with each collapse of the confidence boundary. Note that in the current experiment, with only two levels of confidence, the model simplifies to a single step such that the confidence boundary only takes two heights (i.e. *h* corresponding to high confidence and *h-*∞ corresponding to low confidence).

Third, for the FCB model, two confidence boundaries that both slowly collapse demarcate the area of evidence accumulation. The height of the confidence boundaries are given by equation (3):

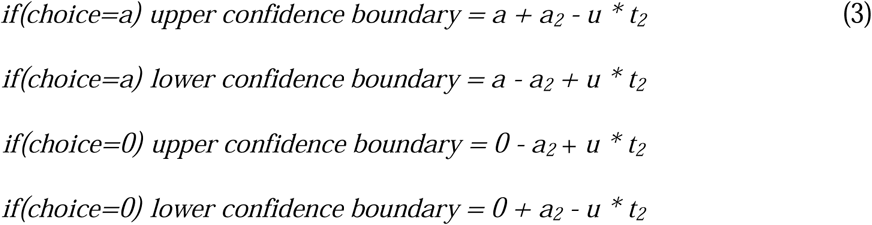

with a_2_ reflecting the height of the confidence boundaries, *u* reflecting the amount of (linear) urgency, and *t_2_*reflecting post-decision time. Given that Experiment 1 only has two levels of confidence (high vs low), confidence here fully coincides with the boundary that was reached (i.e., high versus low confidence when reaching the upper vs lower confidence boundary; see Figure 1D).

Finally, the Van Zandt model was implemented as two independent evidence accumulators which race towards a single decision boundary and then continue to race towards a single confidence boundary with heights *a* and *a_2_*, respectively. The accumulators were defined as specified for the other models, but started at zero and had a separate drift rate for the correct choice accumulator *v_correct_* and the error choice accumulator *v_error_.* RT’s and decision accuracy were defined by the first accumulator to cross the decision boundary. Afterwards, evidence accumulation continued for both accumulators with the same drift rate until one of them crossed the confidence boundary, at which point a confidence judgment was made. If evidence at this point was higher for the first accumulator that reached the decision boundary, a high confidence response was given. Otherwise, the simulation ended with a low confidence response.

Note that in all models an additional time *ter_2_* was added to predicted confidence RTs to capture non-confidence related processes (e.g. pressing a confidence button). In contrast to *ter*, which is by definition always positive, we also allowed *ter_2_* to take negative values, to account for the possibility that post-decision evidence accumulation already starts before an overt response has been made (e.g., during the motor execution of the first response). An overview of the free parameters used in all models can be found in the Supplementary Materials.

### Parameter Estimation and Model Fit

We estimated best fitting parameters separately for each participant and within each condition by jointly minimizing three error functions based on quantile and proportion optimization of the RT and confidence RT distributions, and of the confidence judgment proportions. Quantiles were computed separately for correct and error trials in both observed and simulated data for two types of quantiles, (i) decision reaction time (RT) and (ii) confidence reaction time (RTconf), and the confidence judgment proportions. The resulting error functions are shown in equations (4), (5) and (6):

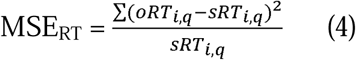

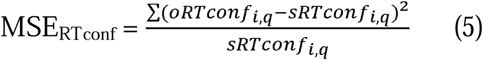

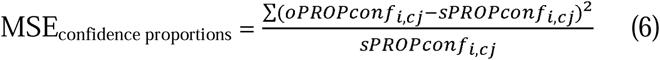

with *oRT* and *sRT* referring to observed and simulated RT proportions, *oRTconf* and *sRTconf* to observed and simulated confidence RT proportions, across multiple quantiles (*q*) (.1, .3, .5, .7, and .9), for correct- and error-trials (*i*) separately. Similarly, oPROPconf and sPROPconf refer to the observed and simulated confidence judgment proportions, across all levels of confidence (*cj*) (0, 1), also for correct- and error-trials (*i*) separately. We minimized the sum of the error functions given by equation (4), (5) and (6) using differential evolution optimization to optimize all parameters. The evolutionary algorithm was operationalized by means of the DEoptim package, with the number of iterations set to 500 and the number of simulated trials set to 2000 (Mullen et al., 2011). Model fitting was done separately per participant. To assess model fits, we simulated choices, RTs, confidence judgments and confidence RTs from the estimated parameters.

Finally, to compare fit across models, we computed BIC values for each fit separately, according to equation (7):

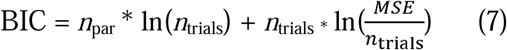

with *n*_par_ and *n*_trials_ referring to the number of free parameters and the number of observed trials, respectively. MSE (Mean Squared Error) indicates the sum of MSE_choice RT_, MSE_confidence RT_ and MSE_confidence proportions_.

## Results

### Behavioral Analysis

Trials with RTs below .2s were excluded from the dataset (00.40%) (Moran et al., 2015). In addition, confidence RTs slower than 5s were excluded (00.10%; note that choice RTs slower than 5s were excluded by design). Next, we report a set of analyses testing how RTs, confidence RTs, accuracy and confidence judgments were influenced by motion coherence (3 levels: .1, .2 and .4), choice SAT (2 levels: fast vs accurate) and confidence SAT (2 levels: fast vs careful).

For reaction times on correct trials (shown in Figure 3A), as expected we found a significant effect of choice SAT instructions, χ^2^(1) = 68.87, *p* < .001, but not of confidence SAT instructions, χ^2^(1) = 0.56, *p* = .455. Choice RTs were shorter when participants were instructed to respond fast (*M* = 0.92s) versus accurate (*M* = 1.38s). Also, the main effect of motion coherence was significant, ^2^(2) = 687.25, *p* < .001, reflecting shorter RTs with increasing motion coherence. Additionally, we found a significant interaction between the choice SAT and confidence SAT, χ^2^(1) = 15.35, *p* < .001, reflecting that the choice SAT effect was more expressed when participants were instructed to provide accurate versus careful confidence ratings. There was also a significant interaction between choice SAT and coherence, χ^2^(1) = 43.24, *p* < .001, reflecting that the choice SAT effect was slightly larger for low coherence trials. All other effects were not significant, *p*s > .525. For accuracy, we likewise found a significant effect of the choice SAT, χ^2^(1) = 8.25, *p* = .004, and coherence, χ^2^(2) = 165.58, *p* < .001, but not of the confidence SAT, χ^2^(1) = 0.63, *p* = .429. As shown in Figure 3B, participants responded more correct when instructed to be accurate (*M* = 72.28%) compared to when instructed to be fast (*M* = 70.78%), and accuracy increased with motion coherence. All other effects were not significant, *p*s > .071.

**Figure 3.**
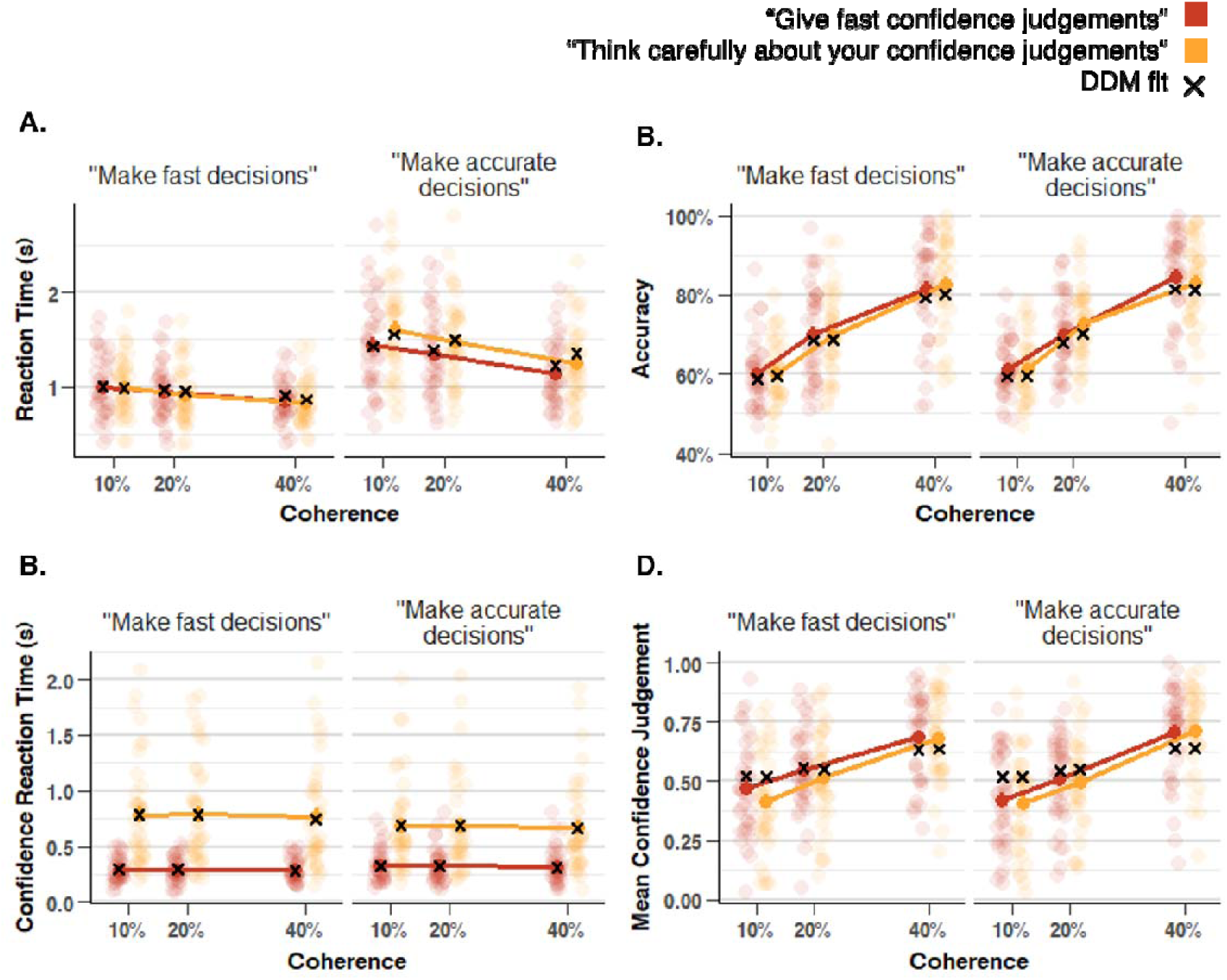
The influence of choice SAT and confidence SAT on reaction times (A), accuracy (B), confidence RTs (C) and confidence (D) for Experiment 1. As expected, when participants were instructed to make fast versus accurate choices this led to fast versus slow choice RTs (A) and to a lesser extent to less and more accurate choices (B), respectively. When participants were instructed to make fast vs deliberate confidence judgments, this led to fast versus slow confidence RTs (C). There was no main effect on confidence judgments (D). Note: error bars reflect SEM, transparent dots reflect means of individual participants, black crosses reflect fits from the FCB model.

For confidence RTs on correct trials, we found significant effects of the confidence SAT instructions, χ^2^(1) = 77.06, *p* < .001, and coherence, χ^2^(2) = 14.29, *p* = .001. As expected, choice SAT instructions did not influence confidence RTs, χ^2^(1) = 0.42, *p* = .518. As can be seen in Figure 3C, confidence RTs were faster when participants were instructed to make fast (*M* = 0.31s) vs careful (*M* = 0.73s) confidence judgments. Additionally, we found a significant interaction between choice SAT and confidence SAT, χ^2^(1) = 6.28, *p* = .012, reflecting a small spill-over from choice SAT into confidence RTs (mostly visible in the “accurate” condition). All other effects were not significant, *p*s > .464. Finally, for confidence judgments (see Figure 3D) we observed a significant main effect of coherence, χ^2^(2) = 120.71, *p* < .001, reflecting that confidence increased with the proportion of motion coherence. There were no significant main effects of choice SAT, χ^2^(1) = 1.23, *p* = .267, nor confidence SAT, χ^2^(1) = 0.27, *p* = .606. There was only a small but significant interaction between choice SAT and confidence SAT, χ^2^(1) = 4.89, *p* = .027, reflecting that participants more often reported high confidence for fast (*M =* .64) than for accurate (*M =* .61) choices in the fast confidence condition, whereas there were was no such difference in the careful confidence condition (.62 vs .62, respectively). Finally, there was an interaction between choice SAT and coherence, χ^2^(2) = 9.97, *p* = .007, reflecting that the relation between confidence and coherence was slightly stronger in the accurate compared to the fast choice condition. All other effects were not significant, *p*s > .160.

Given that there was no effect of the SAT manipulations on average confidence, we additionally examined whether there was a difference in confidence resolution (i.e. the relation between confidence and accuracy). To do so, we ran an exploratory type II ROC analysis separately for each condition (ignoring coherence). A 2-way ANOVA on these estimates showed a main effect of confidence SAT, *F*(1,39) = 15.42, *p* < .001, but not from choice SAT, *p* = .599, nor was there an interaction, *p* = .491. As can be seen in Figure 4A, the relation between confidence and accuracy (expressed in AUC units) was higher when participants were instructed to make deliberate versus fast confidence ratings. Thus, although confidence did not strongly change on average, there was clear evidence that the precision with which participants distinguished correct from incorrect decisions was improved when deliberately computing confidence.

**Figure 4.**
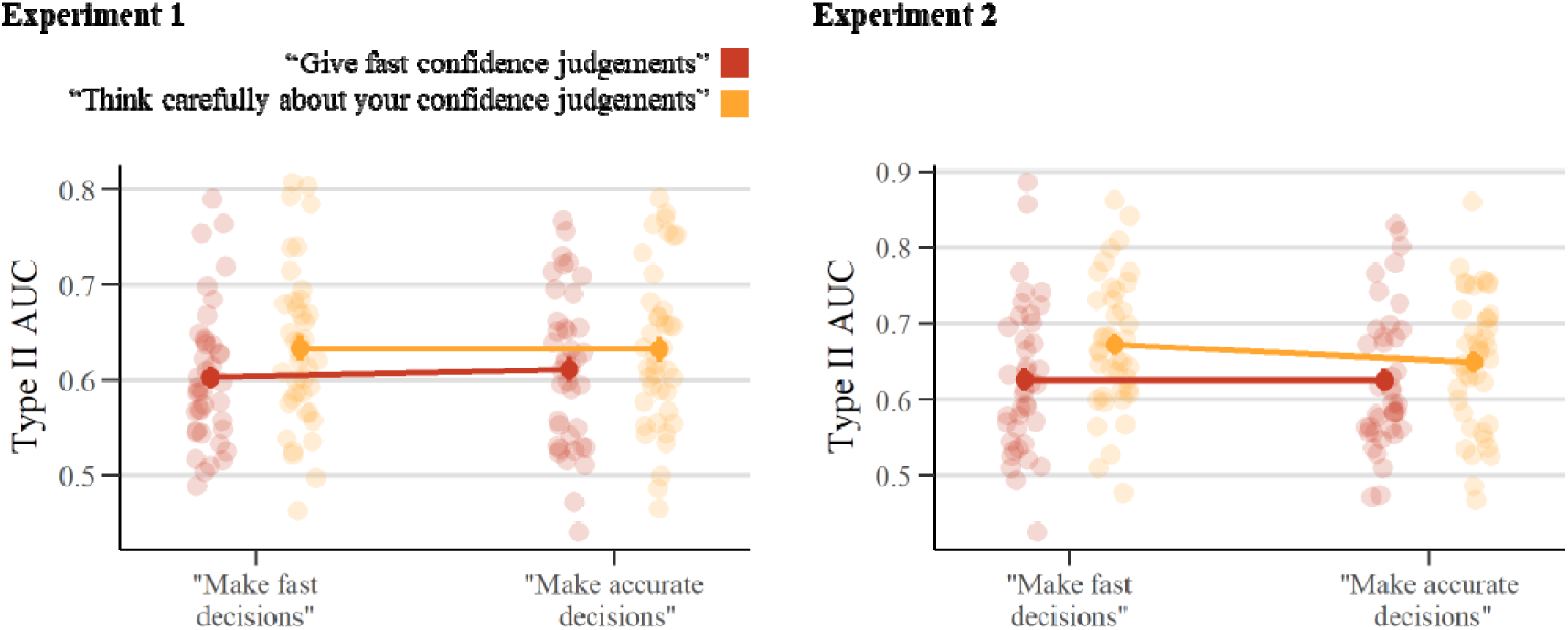
Confidence resolution in Experiment 1 and Experiment 2, expressed as Type II AUC. Although confidence SAT instructions did not have a clear effect on average confidence, we did observe a clear effect on confidence resolution, which was not the case for choice SATs. Same conventions as in Figure 3.

### Modeling the stopping rule for (speed-accuracy tradeoffs in) confidence

Next, we compared the ability of four prominent models of confidence to explain the stopping rule for post-decision evidence accumulation. Our modeling framework departed from the classical drift diffusion model (DDM), a popular evidence accumulation model that accounts well for choices and reaction times in perceptual decisions (Ratcliff & McKoon, 2008). To also account for confidence within the DDM, we allow the evidence to accumulate after it has reached a threshold (post-decision evidence accumulation; Pleskac & Busemeyer, 2010). Critically, we tested different possibilities as to how this process of post-decision evidence accumulation should stop in order to account for the observed data. In the 2DSD, post-decision evidence accumulation terminates after a fixed amount of time has passed. In the CCB model, post-decision evidence accumulation terminates after it reaches a slowly collapsing confidence boundary. In The FCB model, post-decision evidence accumulation terminates once it reaches one of two slowly collapsing confidence boundaries. Finally, according to the model proposed in (Van Zandt & Maldonado-Molina, 2004b), confidence is determined by the difference in evidence between two independent racing accumulators once one of them reaches a (flat) confidence boundary.

#### Model comparison

After fitting the models described above to the data, separately for each participant and condition, we summarized model-averages of MSE and BIC values in Table 1. As can be seen, the FCB model clearly outperforms the competing post-decision evidence accumulation models (ΔBICs > 80). Another thing to note is that the 2DSD model performed particularly worse compared to all three other models (ΔBIC = 2290 compared to the winning model). Although 2DSD performs well in capturing confidence proportions (as shown by the MSE), it completely fails to capture the confidence RT patterns found in the empirical data. The CCB model and the Van Zandt model do a better job compared to the 2DSD in this respect, suggesting that the stopping rule of confidence judgments is best explained by an accumulation-to-bound mechanism and not uniquely by a time-based stopping criterion. Finally, note that the CCB and the Van Zandt model perform worse relative to the FCB model, mostly because the CCB model underperformed both in terms of confidence RT and confidence proportions, and the Van Zandt model captured the observed confidence RTs well, but performed worse in terms of confidence proportions. In sum, results from a model comparison approach show that the stopping rule for confidence is best explained as an accumulation-to-bound process, and specifically as the accumulation towards one of two slowly collapsing confidence boundaries.

**Table 1.** Experiment 1 model comparison. Results from a model comparison shows that the stopping rule for confidence is best accounted for by an accumulation-to-bound mechanism, and specifically by a model (indicated in bold) including post-decision evidence accumulation that terminates ones it reaches one of two opposing slowly collapsing confidence boundaries (i.e. the FCB model).

| Model type | Log RT MSE | Log<br>Confidence<br>RT MSE | Log<br>Confidence<br>proportions<br>MSE | Free<br>parameters | Mean BIC |
| --- | --- | --- | --- | --- | --- |
| 2DSD | 0.47 | 1.70 | -1.26 | 10 | -448.17 |
| CCB | 0.37 | 1.40 | -0.69 | 9 | -518.80 |
| **FCB** | **0.52** | **0.32** | **-1.05** | **9** | **-611.05** |
| Van Zandt | 0.54 | 0.67 | 0.74 | 10 | -531.01 |

Finally we visualized the fit of the winning FCB model. Figures 5A and 5B show that, on average, the predicted confidence RT distributions align well with the observed distributions. Furthermore, the observed patterns of mean RT, confidence RT, accuracy and confidence judgments are captured by the model (see Figure 3). A more in depth analysis of the model fit can be found in supplementary Figure S1.

**Figure 5.**
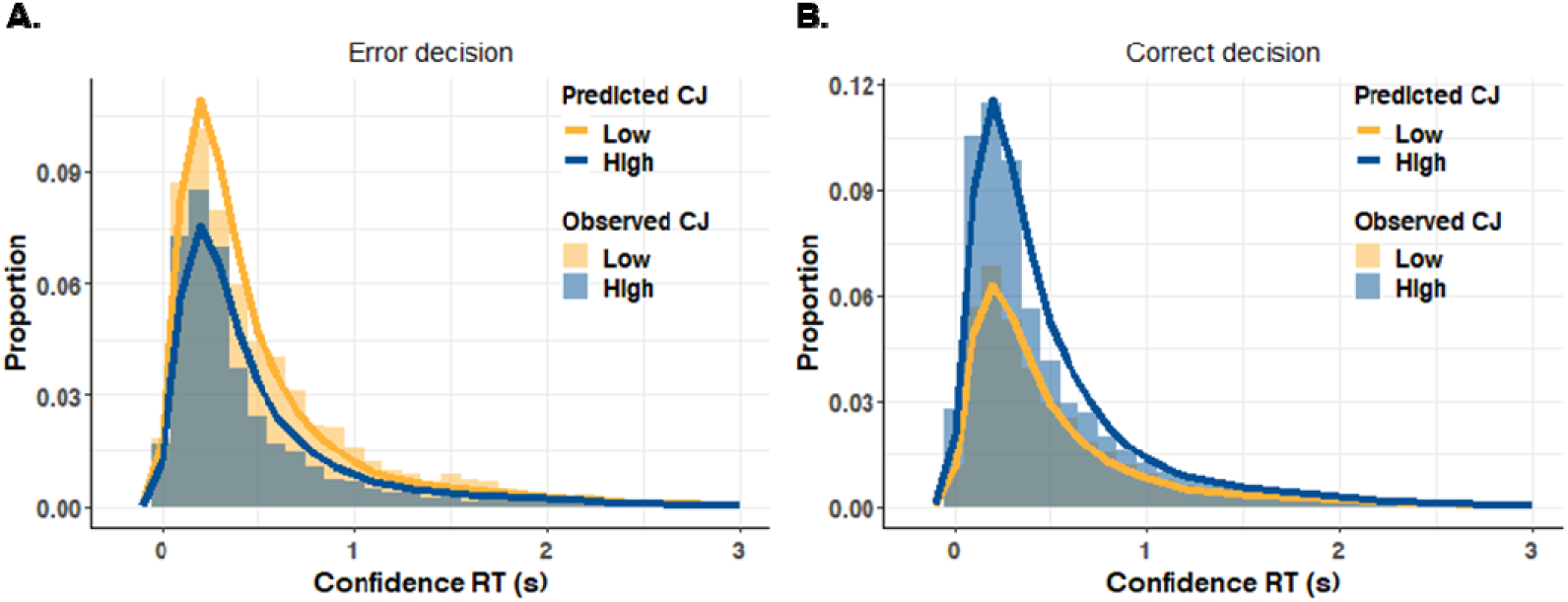
Experiment 1 model fit. Predicted and observed confidence RTs separately for high and low confidence trials after an incorrect (A) and a correct (B) decision by the FCB model. Note, CJ = confidence judgment.

#### Model parameters

Having established that the FCB provides a good fit to the experimental data and outperforms the competing models under consideration, we next turn towards the estimated parameters. We hypothesized that SAT instructions for choices selectively affected choice boundaries, leaving confidence boundaries unaffected. Likewise, we expected SAT instructions about confidence to selectively affect confidence boundaries, leaving choice boundaries unaffected. These observations would support the hypothesis that indeed the stopping rule for both choices and choice confidence are under strategic control. To investigate the interpretability of the model parameters, we first performed a parameter recovery analysis. All parameters recovered well (*r*s > .81), with the exception of the confidence boundary urgency parameter (*u*), which should thus be interpreted with caution. For ease of reading, below we only report the analyses of a priori interest; the entire set of analyses can be found in the supplementary materials (Figures S3, S4).

We used a repeated measures ANOVA to examine the influence of choice SAT (fast vs accurate), confidence SAT (fast vs careful), and their interaction on estimated decision boundaries. As expected, we found a strong and significant effect of the choice SAT, *F*(1, 39) = 70.36, *p* < .001, η_p_^2^ = .64, but no effect of confidence SAT, *F*(1, 39) = 1.50, *p* = .229, η_p_^2^ = .04, nor an interaction between both *F*(1, 39) = 1.98, *p* = .167, η_p_^2^= .05. As can be seen in Figure 6A, when participants were asked to make fast decisions, the separation between both choice boundaries was smaller (*M* = 1.55) compared to when they were asked to make accurate decisions (*M* = 1.99).

**Figure 6.**
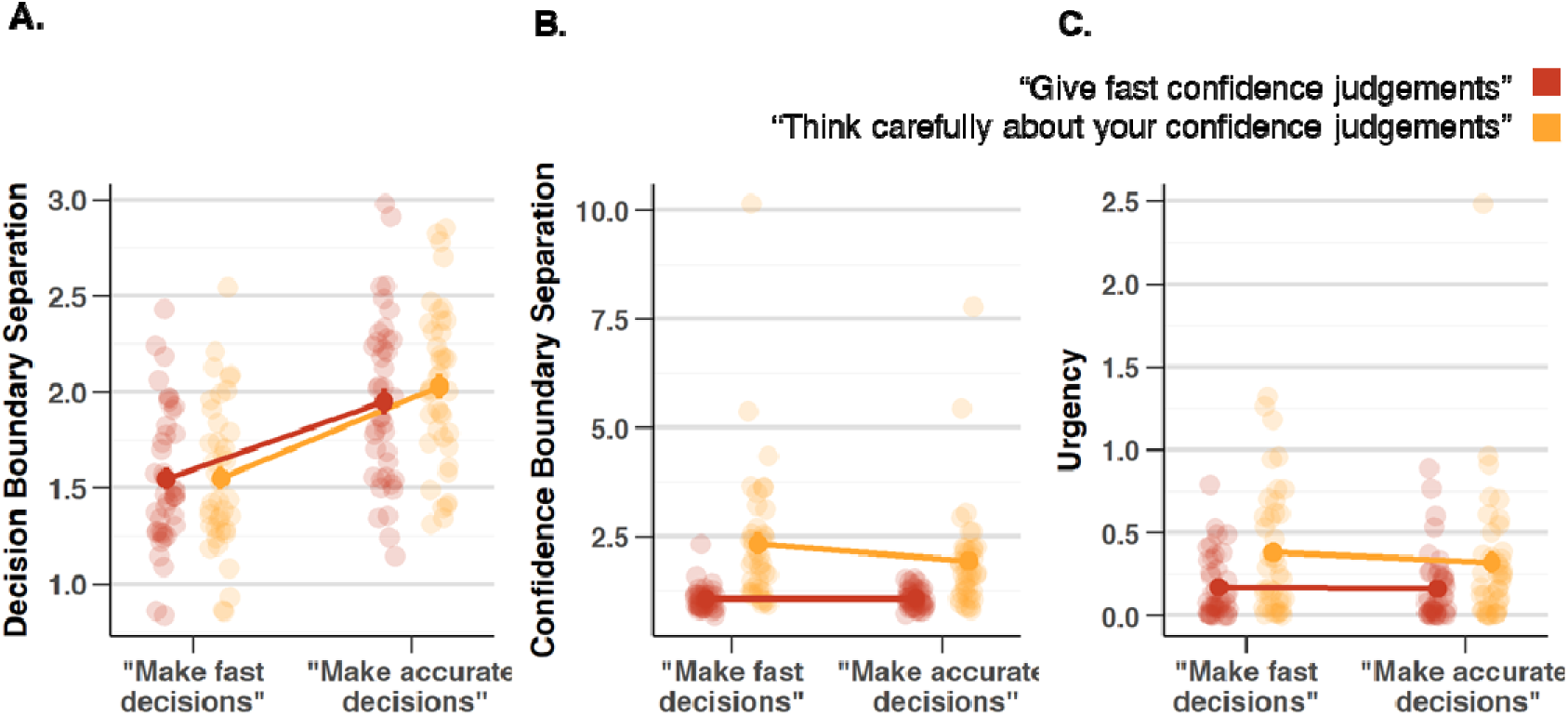
Influence of choice SAT and confidence SAT on decision boundaries and confidence boundaries in Experiment 1. Instructing participants to make fast vs accuracy choices influenced estimated decision boundaries and confidence boundary separation (A, B), but not confidence boundary urgency (C). Instructing participants to provide fast vs careful confidence ratings influenced estimated confidence boundary separation (B) and urgency (C), but did not affect decision boundary separation (A). Same conventions as in Figure 3.

The same analysis on the estimated confidence boundary separation revealed a significant main effect of confidence SAT, *F*(1, 39) = 26.15, *p* < .001, η_p_^2^ = .40, but also of choice SAT, *F*(1, 39) = 8.17, *p* = .007, η_p_^2^ = .17 and the interaction between both types of instruction, *F*(1, 39) = 6.22, *p* = .017, η_p_^2^= .14. Follow-up paired *t*-tests showed that the effect of the confidence RT instructions on confidence boundary separation was significant both when decision SAT was to be accurate, *t*(39) = -4.31, *p* < .001, as well as when decision SAT was to be fast, *t*(39) = -5.14, *p* < .001. As shown in Figure 6B, confidence boundary separation was smaller when participants were asked to make fast confidence judgments (*M* = 1.05) than when participants were asked to make accurate confidence judgments (*M* = 2.13). In sum, choice boundaries were modulated by confidence SAT instructions, but also by decision SAT instructions and the interaction between both.

Third, we looked at the influence of SAT instructions on estimated urgency parameters. Although these should be interpreted with caution, given somewhat lower recovery, the repeated measures ANOVA revealed that there was a significant effect of the confidence SAT, *F*(1, 39) = 17.13, *p* < .001, η_p_^2^ = 0.31, and not of the choice SAT, *F*(1, 39) = 0.58, *p* = .449, η_p_^2^= 0.02, nor was there a significant interaction, *F*(1, 39) = 0.57, *p* = .454, η_p_^2^ = 0.01. Urgency tended to be lower when asked to give fast confidence judgments (*M* = 0.17), compared to when asked to make accurate confidence judgments (*M* = 0.35). Thus, it seems that confidence SAT instructions are implemented by balancing the height and urgency of the confidence boundaries. In Figure 6C, one outlier with an urgency value higher than 2 can be noticed. Removing this outlier from the analysis did not alter any of the conclusions.

Finally, as a sanity check we confirmed that estimated drift rates scaled with motion coherence using a repeated measures ANOVA, *F*(1.28, 49.73) = 173.25, *p* < .001, _ηp_^2^ = .82. The other parameters were not allowed to vary by the instruction conditions, their mean estimates can be found in Table 2.

**Table 2.**
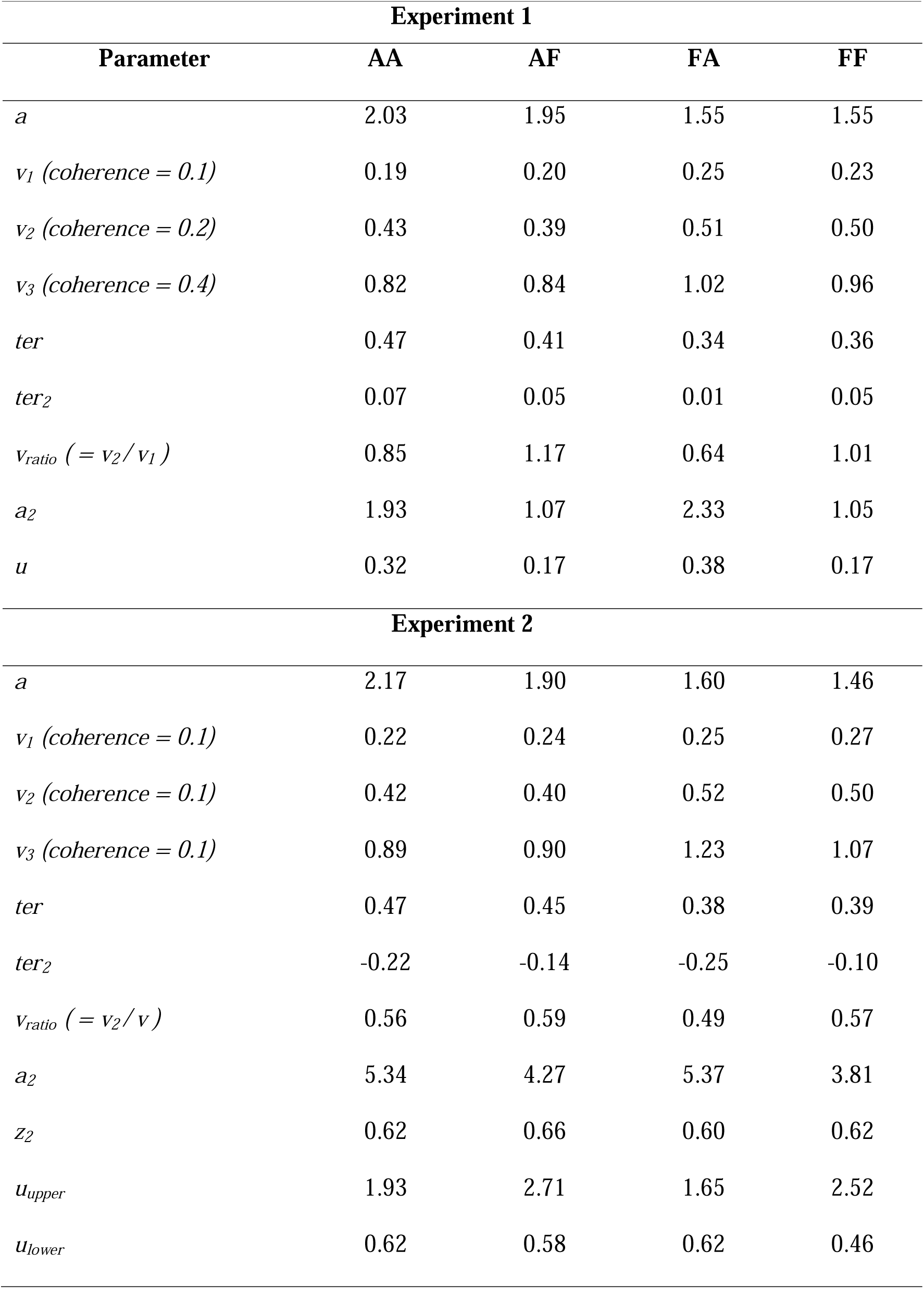
Mean (SD) estimates for the parameters of the best fitting FCB model. Note, AA, AF, FA and FF refer to SAT instructions to be accurate/cautious (A) or fast (F) with the first index referring to the decision and the second index referring to confidence.

### Interim Summary

In Experiment 1, participants were instructed to make fast or accurate decisions and to make fast or careful confidence judgments, depending on the block they were in. At the behavioral level, we observed that participants were indeed able to selectively speed up choices or confidence judgments when instructed to do so. Next, we fitted four different post-decisional evidence accumulation models to explain choices, reaction times, confidence and confidence RTs. Comparison of model fit showed that a DDM with post-decision evidence accumulation until reaching one of two slowly collapsing confidence boundaries, i.e., the newly proposed FCB model, explained the data best. Notably, according to the FCB model, the mechanism underlying speed-accuracy tradeoffs was a modulation of the decision boundary for choices, and a modulation of the confidence boundary for confidence. Thus, these findings suggest that the stopping rule for confidence judgments is best explained as an accumulation-to-bound mechanism, and that just like the choice boundary for choices these confidence boundaries are under voluntary strategic control.

One limitation of Experiment 1 is that participants were only allowed to give binary confidence ratings (high or low). This design choice made for an easy modeling approach, because it allows to directly map high and low confidence onto the upper and lower confidence boundary in the FCB model, respectively. It is well known, however, that humans can provide more fine-grained estimates of their performance. Thus, this begs the question whether the FCB model is also the preferred model in a task with a fine-grained confidence scale. To this end, in Experiment 2 we replicated Experiment 1, but now using a more fine-grained 6-choice confidence scale.

### Experiment 2

#### Methods and Materials

##### Preregistration and Code

The preregistration of this experiment can be found on OSF registries (https://doi.org/10.17605/OSF.IO/VYH4K), all code and data can be found on GitHub (https://github.com/StefHerregods/ConfidenceBounds).

##### Participants

A total of 54 participants participated in Experiment 2. Requirements and recruitment were identical to Experiment 1, with the additional criterium of not having participated in Experiment 1. Data of six participants were removed for not having an accuracy above chance level (as assessed by a binomial test), and four participants for requiring more than seven training blocks. Finally, four participants did not finish the experiment in time. The final sample comprised 40 participants (33 female), with a mean age of 18.5 (*SD* = 1.3, range = 17 - 24).

##### Stimuli and Apparatus

Experiment 2 used the same apparatus and stimuli as in Experiment 1.

##### Procedure

The experiment was identical to Experiment 1, except for the following two exceptions: First, instead of a binary confidence rating, participants could choose between six options; ‘Sure wrong’, ‘Probably wrong’, ‘Guess wrong’, ‘Guess correct’, ‘Probably correct’ and ‘Sure correct’, using the ‘1’, ‘2’, ‘3’, ‘8’, ‘9’ and ‘0’ keys on top of the keyboard. These six options were mapped onto a 1-6 confidence scale (counterbalanced between participants). Second, a time-limit of 5s was imposed on indicating confidence judgments, equal to the time-limit during decision-making. If a participant did not respond within this limit, they were instructed to respond faster in future trials with the following text: ‘Too slow… Please respond faster’.

### Model Specification and fit

Given that participants indicated confidence on a six-point scale, we adapted the models such that they could produce confidence on a similar scale. The 2DSD model now included five confidence criteria (*c*_1_ through *c*_5_; In comparison to only one confidence criterion in Experiment 1) which mapped post-decision evidence onto a six-point confidence scale. The CCB model based on Moran et al. (2015) continued the “staircase-like” pattern reflecting a discrete collapse of a single confidence boundary. As implemented in Moran et al. (2015), the collapse time parameter τ*_cj_*was halved for each successive criterion, and the model produced confidence on a six-point scale. For the FCB model, we changed the implementation such that confidence no longer corresponds to the confidence boundary that was reached (as in Experiment 1). Instead, confidence now depends on the level of accumulated evidence when reaching the confidence boundary. We evenly divided the space in between the two confidence boundaries into six levels, and the model produced a level of confidence between 1 and 6 depending on the state of the accumulated evidence. Note that for Experiment 2 we implemented three additional variants of the FCB model, namely a variant in which we separately estimated urgency for the upper (*u_upper_)* and lower (*u_lower_*) confidence boundary, a model variant in which we allowed the starting point bias for confidence to be freely estimated (*z_2_*), and a model variant in which both of these effects take place. This was done to allow the model to account for various relationships between confidence judgments and confidence RTs (discussed in more detail below). Finally, the Van Zandt race model was adapted to a six-point confidence scale by converting the post-decisional evidence difference between the two accumulator (evidence in favor the chosen option – evidence in favor of the other option) into six categorical values based on the height of the confidence boundary, *a_2_*. Specifically, the range between -*a_2_* and *a_2_* was evenly distributed into six categories, transforming evidence differences to confidence judgments between 1 and 6.

## Results

### Behavioral Analysis: Mixed Effects Modelling

Data were analyzed in the same way as described in Experiment 1. Trials with a decision time of less than 0.2 s were excluded (0.30%). A mixed effects model on choice RTs on correct trials showed a significant effect of choice SAT, χ^2^(1) = 57.91, *p* < .001, and coherence, χ^2^(2) = 956.89, *p* < .001. Unexpectedly, there also was a significant effect of confidence SAT, χ^2^(1) = 34.02, *p* < .001. Additionally, we found significant interactions between the choice SAT and confidence SAT, χ^2^(1) = 8.72, *p* = .003, between coherence and choice SAT, χ^2^(2) = 21.80, *p* < .001, and between coherence and confidence SAT, χ^2^(2) = 12.10, *p* = .002. The three-way interaction between choice SAT, confidence SAT and coherence was not significant, χ^2^(2) = 0.65, *p* = .722. As can be seen in Figure 7A, choice RTs were shorter when participants were instructed to respond fast (*M* = 0.93s) versus accurate (*M* = 1.34s), however the effect was not as selective as in Experiment 1, because choice RTs were also shorter when participants were instructed to provide fast (*M* = 1.06s) versus careful confidence ratings (*M* = 1.20s). The same analysis on accuracy likewise showed significant main effects of choice SAT, χ^2^(1) = 6.05, *p* = .014, confidence SAT, χ^2^(1) = 4.76, *p* = .029, and coherence χ^2^(2) = 1202.14, *p* < .001 (see Figure 7B). Accuracy was lower when participants were instructed to make fast (*M* = 73%) compared to accurate choices (*M* = 75%), and likewise when participants were instructed to make fast (*M* = 74%) versus careful confidence ratings (*M* = 75%). All other effects were not significant, *p*s > .257.

**Figure 7.**
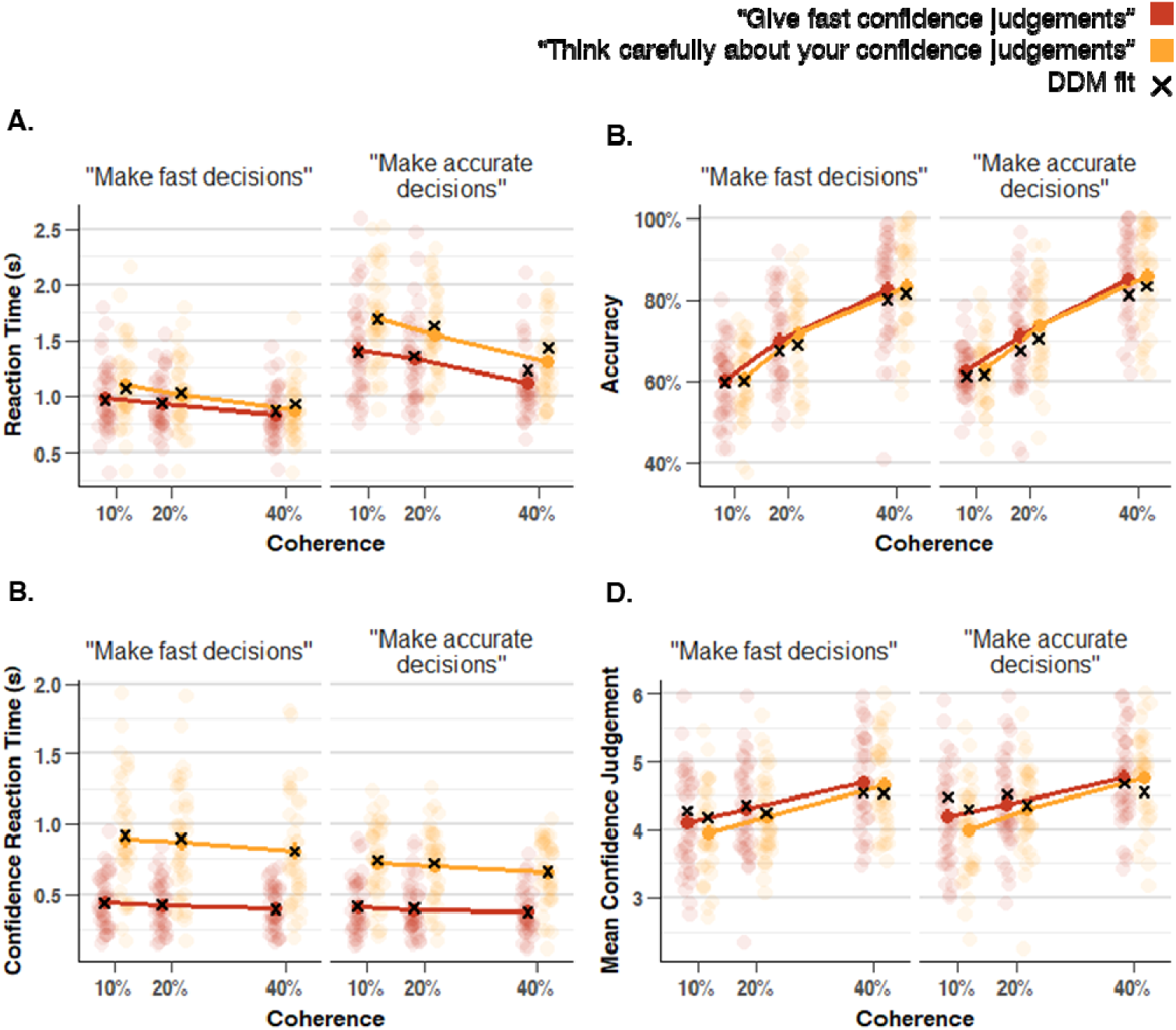
The influence of choice SAT and confidence SAT on reaction times (A), accuracy (B), confidence RTs (C) and confidence (D) for Experiment 2. As expected, when participants were instructed to make fast versus accurate choices this led to fast versus slow choice RTs (A) and to a lesser extent to less and more accurate choices (B), respectively. When participants were instructed to make fast vs deliberate confidence judgments, this led to fast versus slow confidence RTs (C), with less pronounced effects on confidence judgments. Note: same conventions as in Figure 3.

The same analysis on confidence RTs on correct trials, showed significant main effects of confidence SAT, χ^2^(1) = 85.62, *p* < .001, and coherence, χ^2^(2) = 71.12, *p* < .001. Unexpectedly, there was also a significant main effect of choice SAT, χ^2^(1) = 9.64, *p* = .002. Finally, the interaction between the confidence SAT and coherence was significant, χ^2^(2) = 6.26, *p* = .044. All other effects were not significant, *p*s > .161. As can be seen in Figure 7C, although confidence SAT clearly affected confidence RTs in the expected way, the effect was not as selective as in Experiment 1. Confidence RTs were shorter when participants were instructed to make fast (*M* = .39s) versus careful (*M* = .76s) confidence ratings, and counterintuitively confidence RTs were slightly longer when participants were instructed to make fast (*M* = .62s) versus accurate (*M* = .53s) decisions.

Finally, the same analysis was carried out on confidence for correct trials. Note that for this analysis, the three-way interaction and the interaction between the choice SAT and confidence SAT were excluded because they caused variance inflation factors higher than 10. In the final model, there was a significant main effect of coherence, χ^2^(2) = 100.06, *p* < .001, and the confidence SAT, χ^2^(1) = 4.36, *p* = .037, but not of the choice SAT, χ^2^(1) = 0.84, *p* = .359. As can be seen in Figure 7D, variations in confidence were mostly driven by coherence, but confidence was also slightly lower when participants were instructed to make fast (*M* = 4.54) versus careful (*M* = 4.84) confidence judgments. Finally, we found a significant interaction between the confidence SAT and coherence, χ^2^(2) = 22.10, *p* < .001, reflecting that the confidence SAT was more pronounced on low coherence trials. The interaction between the choice SAT and coherence was found to be not significant, χ^2^(2) = 1.57, *p* = .455.

Similar to Experiment 1, in a non-preregistered analysis we additionally looked at confidence resolution by calculating type II AUC separately for each condition. Again, a 2-way ANOVA showed a main effect of confidence SAT, *F*(1,39) = 14.49, *p* < .001, but not from choice SAT, *p* = .066, nor was there an interaction, *p* = .125. As can be seen in Figure 4B, the relation between confidence and accuracy (expressed in AUC units) was higher when participants were instructed to make careful vs fast confidence ratings, suggesting that participants gave more accurate confidence ratings in the careful condition.

### Modelling speed-accuracy trade-offs in 6-point scale confidence judgments

#### Model comparison

We again fitted each model to the data of all 40 participants and computed BIC values. In addition to fitting the 6-choice versions of the models also used in Experiment 1 (2DSD, CCB, FCB and Van Zandt), we additionally included three different variants of the FCB model: FCB_seperate u2_, a variant with separate urgencies for the upper and low confidence boundary, FCB_free z2_, a variant with the starting point for post-decision accumulation as a free parameter, and FCB_seperate u2 and free z2_, a variant with both separate urgency parameters for the confidence boundaries and a free parameter for the post-decision starting point.

Results from the model comparison are shown in Table 3. In line with the findings of Experiment 1, three out of four variants of the FCB model outperform the other models (ΔBICs > 14.28). Within the FCB family, the variant with separate urgency parameters for the confidence boundaries and a free starting point for post-decision accumulation , FCB_seperate u2 and free z2_, provided the best fit to the data (ΔBICs > 60.82). To further unravel why models of the FCB family provide a better fit compared to the competing models, we investigated which specific features could not be accounted for by these models. In line with the findings of Experiment 1, the 2DSD and CCB models fail to capture some of the confidence RT patterns, because of the shape of their stopping criteria. To visually represent this, Figure 8A shows the relation between confidence and confidence RTs for a representative example participants. As can be seen, while this participant showed an inverted-U association between confidence and confidence RTs, the 2DSD model predicts the same confidence RT for all levels of confidence, whereas the CCB model predicts a negative confidence RT – confidence judgment relation. The Van Zandt race model fits confidence RTs relatively well, but it still provides an overall worse fit because it does not capture the observed confidence proportions well (i.e. as shown in Table 3). The best model fit to our data was FCB_seperate u2 and free z2_, in which confidence boundaries had separate urgency parameters and the post-decision starting point could freely vary. As shown in Figure 8A, this model was able to account for the inverted-U shaped association between confidence and confidence RTs seen for this representative example participant.

**Figure 8.**
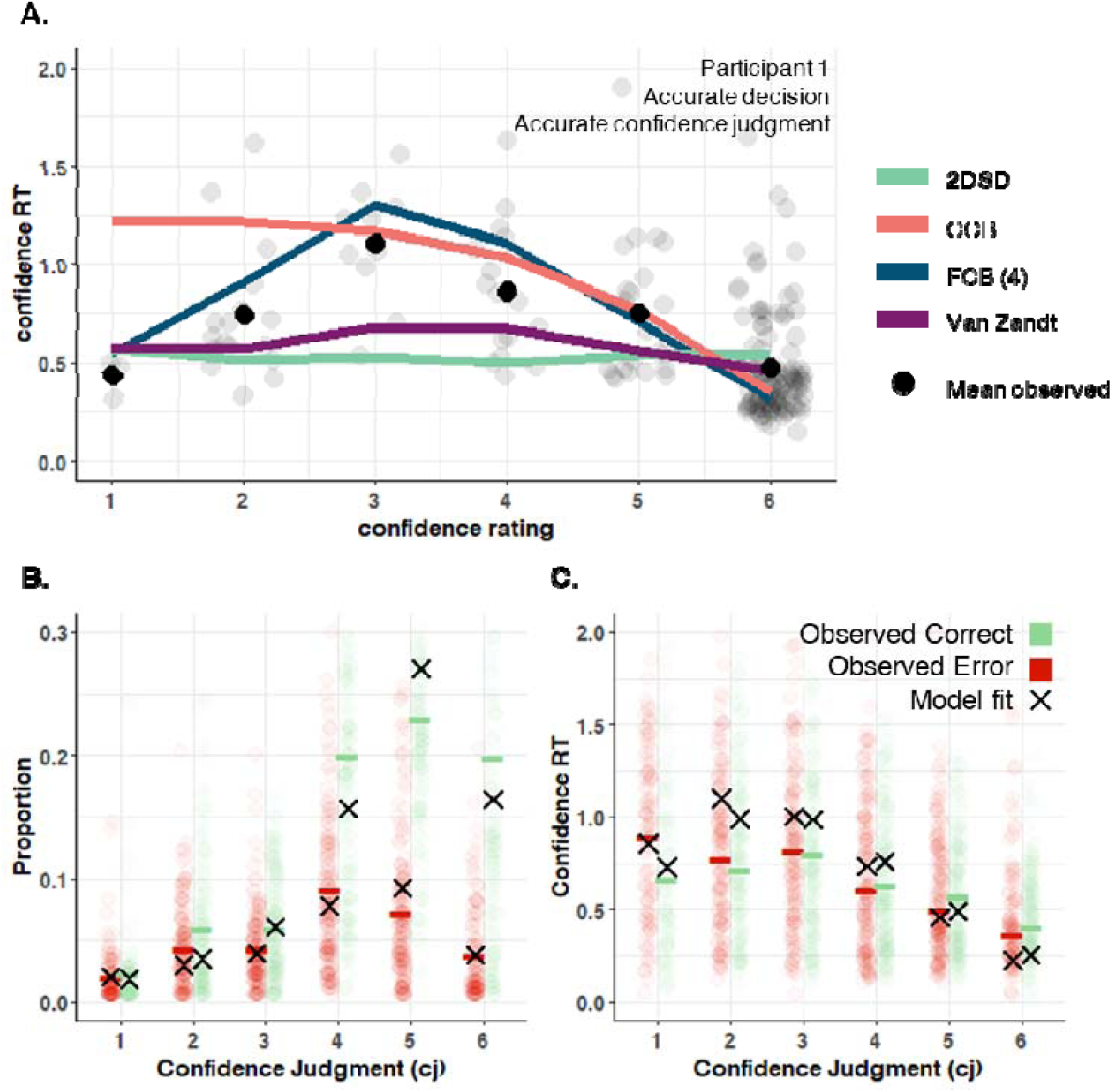
Experiment 2 model fit. **A.** The relation between confidence and confidence RTs for a representative example participant, together with predictions of the different models. As can be seen, only the FCB_separate u2 and free z2_ (and to some extent the Van Zandt) model is able to capture the inverted U-shape relation between confidence RT and confidence ratings. , **B-C**. Observed versus predicted confidence proportions (B) and RTs (C) by the best performing model (FCB_seperate u2 and free z2_)).

**Table 3.**
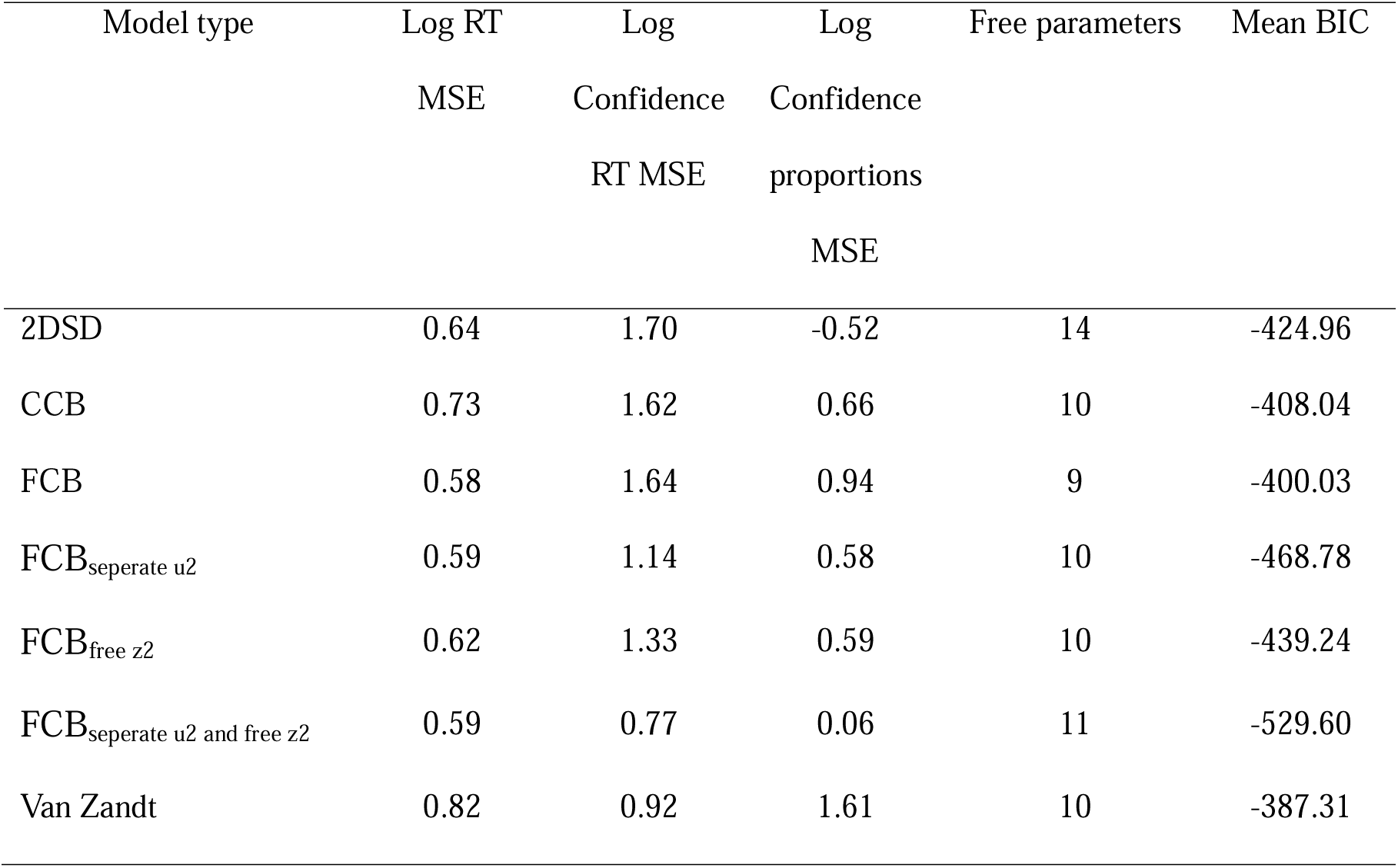
Experiment 2 model comparison.

Finally, In Figure 8B-C it can be seen that the best performing model, FCB_seperate u2 and free z2_, closely captured the observed confidence proportions (Figure 8B), and confidence RTs (Figure 8C). Additionally, the model captured the RT, confidence RT, accuracy and confidence judgment patterns described in Figure 7. A thorough examination of the model fit is available in Supplementary Figure S2.

#### Model parameters

Next, we investigated the influence of the SAT instructions on the parameters of the best fitting model, the FCB model with separate urgencies and a flexible post-decision starting point. A parameter recovery analysis of this model showed that all parameters recovered well, including the confidence urgency parameters (which were difficult to recover in the case of binary confidence judgments). Full results of this analysis can be found in the supplementary materials (Figures S5, S6).

Similar to the findings of Experiment 1, we again observed that estimated decision boundaries were affected by choice SAT, *F*(1, 39) = 66.41, *p* < .001, η_p_^2^ = .63. However, we also found a significant effect of the confidence SAT, *F*(1, 39) = 31.90, *p* < .001, η_p_^2^ = .45, and an interaction between both, *F*(1, 39) = 4.82, *p* = .034, η_p_^2^ = .11. Follow-up paired *t*-tests showed that the effect of the choice RT instructions on decision boundary separation was significant both when confidence SAT was to be accurate (*M* = 1.89), *t*(39) = -7.75, *p* < .001, as well as when confidence SAT was to be fast (*M* = 1.68), *t*(39) = -6.87, *p* < .001. As expected, choice boundaries were modulated by choice SAT instructions (Figure 9A), although the effect also seemed to scale, to a lesser extent, with confidence SAT.

**Figure 9.**
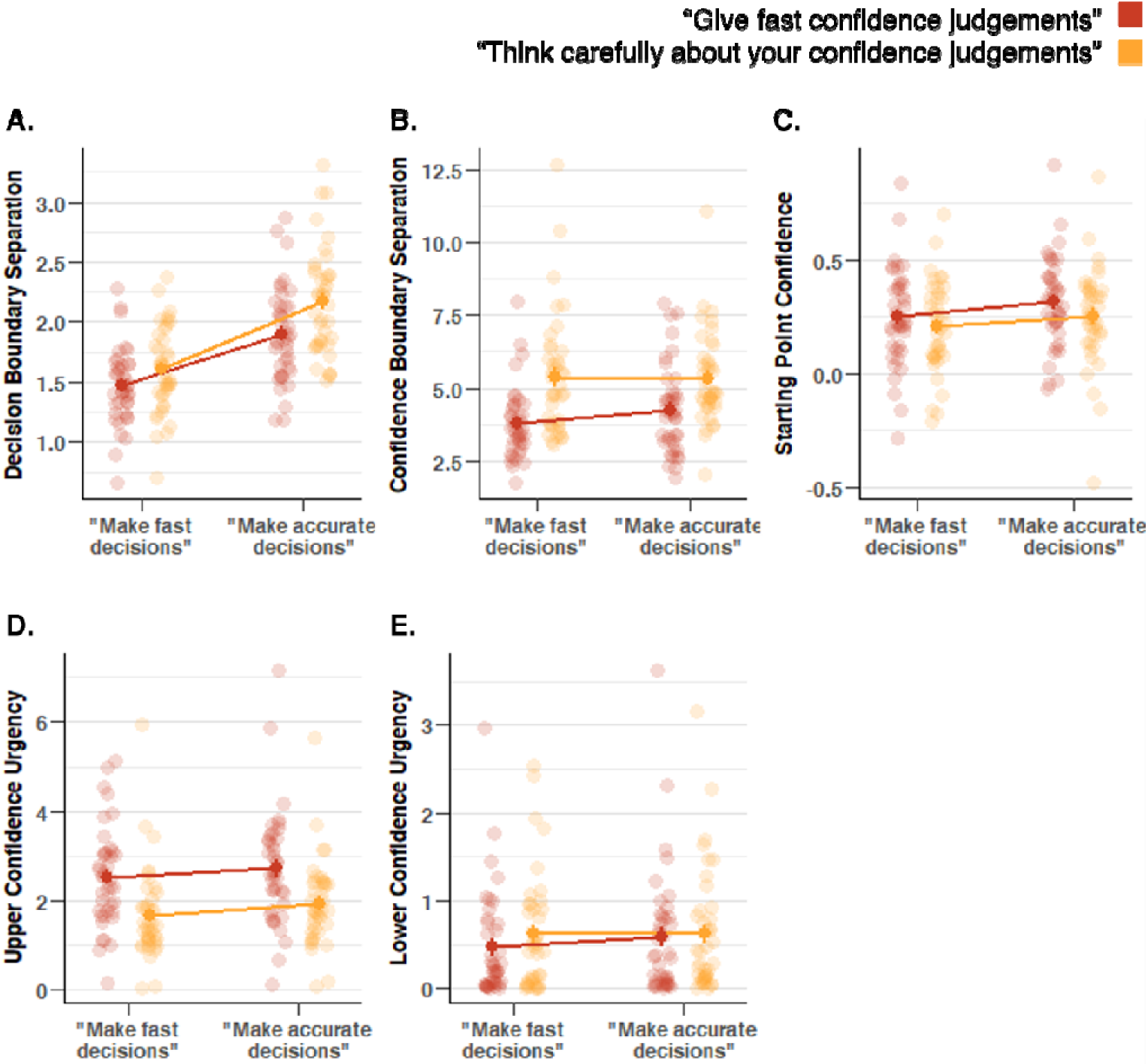
Influence of choice and confidence SAT on decision boundaries and confidence boundaries in Experiment 2. Instructing participants to make fast vs accuracy choices influenced estimated decision boundaries (A) and the starting point of post-decision evidence accumulation (C), but not confidence boundary separation or upper/lower confidence urgency (B, D, E). Instructing participants to provide fast vs careful confidence ratings influenced the decision boundary separation, starting point of post-decision evidence accumulation, confidence boundary separation and upper confidence boundary urgency (A-D), but not lower confidence boundary urgency (E). Same conventions as in Figure 3.

Second, we analyzed the confidence boundary separation. In line with findings from Experiment 1, we found a significant effect of confidence SAT, *F*(1, 39) = 58.42, *p* < .001, η_p_^2^ = .60. However, we did not find a significant effect of decision SAT, *F*(1, 39) = 1.35, *p* = .252, η_p_^2^ = .03, or the interaction between both, *F*(1, 39) = 2.14, *p* = .151, η_p_^2^ = .05. Participants increased confidence boundary separation when instructed to make careful confidence judgments (*M* = 5.36) compared to when instructed to make fast confidence judgments (*M* = 4.04, Figure 9B). Furthermore, we found a significant effect on the starting point of evidence accumulation for confidence of confidence SAT, *F*(1, 39) = 29.79, *p* < .001, η_p_^2^= .43, and of decision SAT, *F*(1, 39) = 14.37, *p* < .001, η_p_^2^ = .27, but not of the interaction between both, *F*(1, 39) = 1.04, *p* = .315, η_p_^2^ = .03. More specifically, participants tended to start evidence accumulation for confidence judgments at a higher level of evidence when asked to give fast confidence judgments (*M* = 0.64), than when asked to think more carefully about them (*M* = 0.61), but the starting point was also influences by the decision SAT instructions (Figure 9C).

Notice that, different from Experiment 1, both confidence boundary urgencies were allowed to vary independently and thus are analyzed separately. Analysis of the upper confidence boundary urgency revealed a significant effect of the confidence SAT, *F*(1, 39) = 38.98, *p* < .001, η_p_^2^ = .50. We did not observe an effect of the choice SAT, *F*(1, 39) = 3.66, *p* = .063, η_p_^2^ = .09, nor the interaction between both, *F*(1, 39) = 0.18, *p* = .674, η_p_^2^ = .01. As shown in Figure 9D, the slope was steeper when participants were instructed to make fast confidence judgments (*M* = 2.61), than when given the instruction to think carefully about their confidence judgments (*M* = 1.79). Finally, for the lower confidence boundary we found that urgency was not affected by confidence SAT, *F*(1, 39) = 1.17, *p* = .285, η_p_^2^ = .03, choice SAT, *F*(1, 39) = 0.20, *p* = .658, η_p_^2^ = .005, nor was there an interaction, *F*(1, 39) = 0.66, *p* = .422, η_p_^2^ = .02 (Figure 9E).

As a final sanity check, we again confirmed that estimated drift rate scaled with motion coherence, *F*(1.14, 44.30) = 127.25, *p* < .001, η_p_^2^ = .77. All parameter estimates can be found in Table 2.

## Discussion

The human ability to estimate and report the level of confidence in their decisions has been the central topic of many recent investigations (Rahnev et al., 2022). Despite a large number of studies examining how confidence is computed, the question of how people decide *when* to provide a confidence rating has been unresolved. This is remarkable, because the timing of confidence judgments can be highly diagnostic about the computations underlying decision confidence (Moran et al., 2015). In the current work, we propose to model the stopping rule for confidence judgments using an accumulation-to-bound mechanism similar to the one underlying decisions (i.e. the Flexible Confidence Boundary model; FCB), and compare it to three other post-decisional evidence accumulation models, the CCB proposed by Moran et al. (2015), 2DSD proposed by Pleskac and Busemeyer (2010) and the race model proposed by Van Zandt and Maldonado-Molina (2004). We manipulated the stopping rule for confidence judgments by providing participants with different instructions regarding the tradeoff between speed and accuracy, both for decisions and for confidence judgments. In two experiments, we found that participants made faster and less accurate decisions when instructed to favor speed over accuracy, and that they made faster confidence judgments when instructed to favor speed over careful deliberation. Although the effects on average confidence were subtle or even absent, in both experiments the relation between confidence and accuracy (cf. confidence resolution) was clearly stronger when participants were more cautious in their confidence ratings. Most importantly, across both experiments we found through qualitative and quantitative model comparison that the newly proposed FCB model fitted the data better than the alternative models. Inspection of the estimated parameters of winning FCB model showed that, as expected SAT instructions about the decision influenced decision boundaries, and SAT instructions about confidence influenced confidence boundaries. Our findings have important consequences for the field of decision confidence, as they shed light on the stopping rule for confidence and thereby unravel the importance of considering the dynamics of confidence RTs.

### Modelling the stopping rule of evidence accumulation for confidence

Previous work investigating the dynamics of decision confidence has mostly focused on explaining variations in decision confidence, with less focus on the speed with which confidence reports are given. If confidence RTs are considered during modeling, the most common approach is to simply have a free parameter that controls the duration of post-decision evidence accumulation (Hellmann et al., 2021; Pleskac & Busemeyer, 2010; Yu et al., 2015). This implementation, however, predicts that confidence judgments will always be provided at the same post-decision latency. This prediction is at odds with the observation that confidence RTs show a right-skewed distribution that is also characteristic of choice RTs. Confidence boundaries, on the other hand, provide a plausible mechanism for the stopping rule of post-decision evidence accumulation. Two notable exception to this critique are the model CCB model proposed by Moran and colleagues (2015) and the race model proposed by Van Zandt and Maldonado-Molina (2004). Moran and colleagues proposed a single confidence boundary that collapses slowly over time, with the level of confidence being determined by the height of the boundary at the time of crossing. The proposal from Moran and colleagues differs from our newly proposed FBC model because it does not consider a lower confidence boundary; the model provides a confidence rating of .5 if the collapsing confidence boundary has not been reached before it collapsed to .5. Contrastingly, the FCB model features both an upper and a lower confidence boundary, which can be mapped onto high versus low confidence (Experiment 1), or accounts for graded levels of confidence by further dividing the area in between the two confidence boundaries (Experiment 2). Finally, (Van Zandt & Maldonado-Molina, 2004) proposed that confidence can be computed based on the difference in post-decisional evidence between two independent (racing) accumulators. According to this model, a confidence judgment is given once one of the accumulators crosses a confidence boundary. Results from our model comparison found unequivocal support, across two experiments, that the stopping rule for confidence judgments is best explained by an accumulation-to-bound mechanism. In both experiments, the 2DSD model provided a considerably worse fit to the confidence RT data, precisely because it was unable to capture intricate associations between confidence and confidence RTs. Additionally, from the different models considered here that did implement an accumulation-to-bound mechanism as the stopping rule for confidence, the newly proposed FCB model provided the best fit to the data. Again, the reason why this model was favored over the others was that it was able to flexibly allow for both positive, absent, negative and even U-shaped associations between confidence and confidence RT.

Interestingly, there has been a large body of work that modelled the stopping rule of evidence accumulation in tasks with simultaneous decision-making and confidence judgment tasks (Ratcliff & Starns, 2009, 2013). Given the simultaneous reporting of choice and confidence a single boundary typically suffices to explain choice and confidence latencies, and this line of work proposes multiple accumulators to jointly explain choices and confidence. Such models can effectively account for the timing of choices and confidence, however it does not consider temporal dissociations between choice and confidence. Our work differs substantially from this approach, as our main aim is to explain behavior in tasks where choices and confidence judgments are given sequentially, thus allowing for the accumulation of post-decision evidence.

### The stopping rule for confidence is under strategic control

Speed-accuracy tradeoffs can be implemented in accumulation-to-bound models via two different mechanisms: changing the overall height of the boundary or changing the collapse rate of the boundary over time. Previous work in decision making has unraveled that instructions regarding the tradeoff between speed and accuracy for the decision tend to modulate the height of the decision boundary, while not affecting urgency (Katsimpokis et al., 2020). Reversely, when providing participants with a response deadline (e.g. respond within 1s) data are best accounted for by a slowly collapsing decision boundary (Katsimpokis et al., 2020; Murphy et al., 2016). In theory, the same two mechanisms could be used to implement speed-accuracy tradeoffs for confidence judgments. However, in our experiments where confidence SAT was modulated by means of instructions, given the evidence cited above, it was expected that participants would change the height of the confidence boundary, while leaving the urgency constant. In both our experiments, there was clear evidence that participants changed the height of the confidence boundary in response to SAT instructions. Results were more mixed concerning urgency. In Experiment 1, we observed a significant difference between conditions in terms of urgency, suggesting that participants selectively changed the height of the confidence boundaries in combination with the slope. In Experiment 2, we found that in response to instructions requiring fast confidence responses, participants increased the level of urgency for the upper confidence boundary, but not the lower confidence boundary. Given that both experiments were identical to each other except for the number of confidence options, this suggest that complexity of the design (i.e. arbitrating between high and low confidence versus arbitrating between six fine-grained levels of confidence) is an important factor determining the specificity with which these manipulations have an effect. As already noted, however, parameter recovery for urgency in Experiment 1 was rather low, suggesting these values should be interpreted with great caution. Nevertheless, in addition to further unravelling the role of design complexity, future work might also investigate the influence of providing a hard deadline for confidence judgments (e.g. you have to provide a confidence rating within 1s) on the confidence boundary and associated urgency signal.

### Characteristics of post-decision processing

If confidence can be understood as an accumulation-to-bound signal, it follows that the reported level of confidence should depend on the height of the confidence boundary. Similar to how decreasing the decision boundary induces faster RTs and less accurate responses, it follows that, given a positive drift rate, decreasing the confidence boundaries should induce faster confidence RTs and lower confidence. Contrary to this, in the estimated parameters of the FCB model, we did not observe a clear influence of confidence SAT on average confidence despite a clear difference in the height of the confidence boundary. As can be seen in Table 2, the FCB model explained these data by assuming that decreasing the confidence boundaries was associated with increased post-decision drift rates. Future work might examine whether this prediction holds in post-decision centro-parietal EEG signals, which are thought to reflect the post-decision accumulation-to-bound signal (Desender et al., 2021). Although we did not find an effect on average confidence, there was a clear effect on confidence resolution: the relation between confidence and accuracy was much stronger when participants increased the confidence boundary. This finding could be anticipated, because increasing the confidence boundaries effectively requires collecting more post-decision evidence before reporting confidence, i.e. making a more informed confidence judgment. This finding adds to a number of reports showing that measures of metacognitive accuracy critically depend on the timing of confidence reports (Rosenbaum et al., 2022; Yu et al., 2015).

Inspection of the estimated FCB model parameters in Table 2 reveals an interesting difference in magnitude between the non-decision component associated with the decision, *Ter*, and that associated with the confidence report, *Ter_2_*. In line with the literature, values of *Ter* are in the range of .4s - .5s on average, suggesting that this is the time participants spend on processes unrelated to the actual decision (e.g. stimulus processing, motor components). These estimates are by definition positive. Contrary to this, values of *Ter_2_*are very low for Experiment 1 and even negative for Experiment 2. Although negative values of *Ter_2_* might seem counterintuitive at first, they suggest that “post-decision” processing already initiates prior to the execution of the decision motor response (e.g., Verdonck et al., 2020). In line with this observation, there is some work that has suggested that pre-choice or peri-choice neural signals contribute to the computation of decision confidence (Feuerriegel et al., 2022; Gherman & Philiastides, 2015; Murphy et al., 2015).

## Conclusion

We demonstrated that the stopping rule for confidence judgments is well described by an accumulation-to-bound process terminating post-decision processing, similar to how choice formation occurs. Similar to the decision boundaries, these confidence boundaries are under strategic control, and can be increased or decreased by instructing participants to make very careful or very fast confidence judgments, respectively. This implementation of the stopping rule fits the data better compared to the popular 2DSD model (in which post-decision processing terminates after a fixed amount of time has passed) and two other competing models. Taken together, the current work unravels the stopping rule for post-decision processing that informs the computation of confidence.

## Acknowledgments

This research was supported by a project grant by the Research Foundation Flanders, Belgium (FWO-Vlaanderen No. G0B0521N) a starting grant from the KU Leuven (STG/20/006) and the Francqui Foundation (PXF-D8830), a CELSA grant from the KU Leuven (CELSA/21/010) and a FWO postdoctoral fellowship by the Research Foundation Flanders, Belgium (FWO-Vlaanderen No. 1242924N).

## Supplementary Materials

**Table S1.** Free parameters of the 2DSD model.

| Parameter | Meaning | Description |
| --- | --- | --- |
| $a$ | Decision boundary separation | Distance between decision boundaries |
| $v$ | Drift rate | Average increase of evidence towards the correct decision boundary |
| $ter$ | Non-decision time | Time during the decision-making process where evidence is not being accumulated (e.g., the time between deciding which button to press and the actual button press because of motor execution time) |
| $\tau_{cj}$ | Mean interjudgment time | Average post-decision evidence accumulation time across trials |
| $SD\tau$ | Standard deviation<br>Interjudgment time | Standard deviation of post-decision evidence accumulation time across trials |
| $c_1$ | First confidence threshold | Criterion indicating the boundary of post-decision evidence between the lowest and second-lowest confidence judgment. Defined relative to the amount of evidence accumulated when crossing the decision boundary. |
| $c_2, c_3, c_4,$<br>$c_5$ | Confidence thresholds | Confidence thresholds indicating the boundary of post-decision evidence between higher confidence judgments. Defined relative to the |
|  |  | previous confidence threshold. Only used for Experiment 2. |
| $v_{ratio}$ | $v$-ratio | Ratio between the pre- and post-decision drift rate |
| $ter_2$ | Confidence non-decision time | Time during the process of deciding on a confidence level where evidence is not being accumulated |

**Table S2.** Free parameters of the CCB model.

| Parameter | Meaning | Description |
| --- | --- | --- |
| $a$ | Decision boundary separation | Distance between decision boundaries |
| $v$ | Drift rate | Average increase of evidence towards the correct decision boundary |
| $ter$ | Non-decision time | Time during the decision-making process where evidence is not being accumulated |
| $h$ | First confidence boundary height | Absolute height of the highest confidence boundary, relative to the amount of evidence accumulated when crossing the decision boundary. |
| $\Delta$ | Collapse height | Collapse height of the confidence boundaries once the collapse time is reached |
| $\tau$ | First collapse time | Time before the first confidence boundary |
|  |  | collapses. Each consecutive collapse time is half of the previous collapse time. |
| $v_{ratio}$ | $v$-ratio | Ratio between the pre- and post-decision drift rate |
| $ter_2$ | Confidence non-decision time | Time during the process of deciding on a confidence level where evidence is not being accumulated |

**Table S3.**
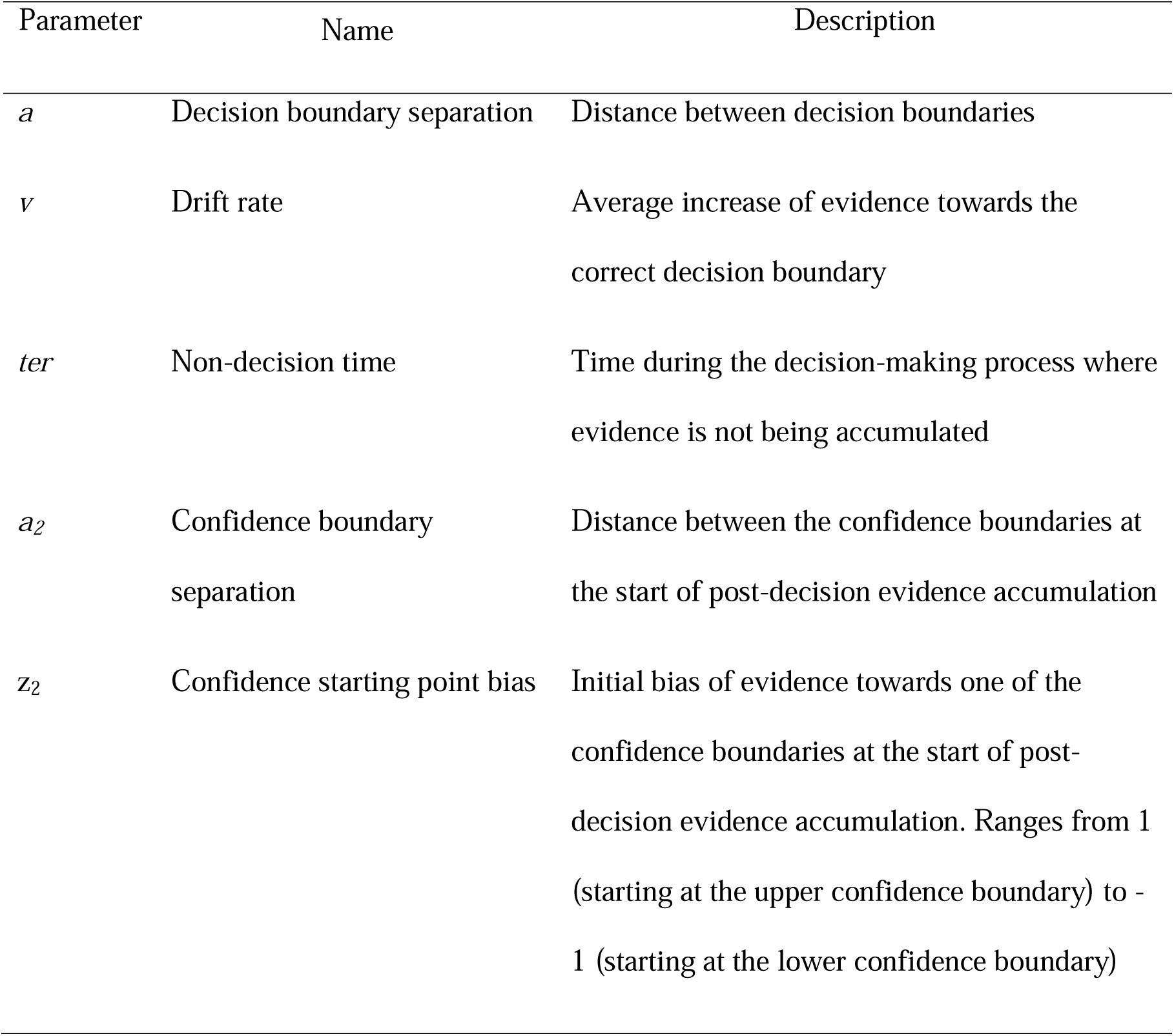
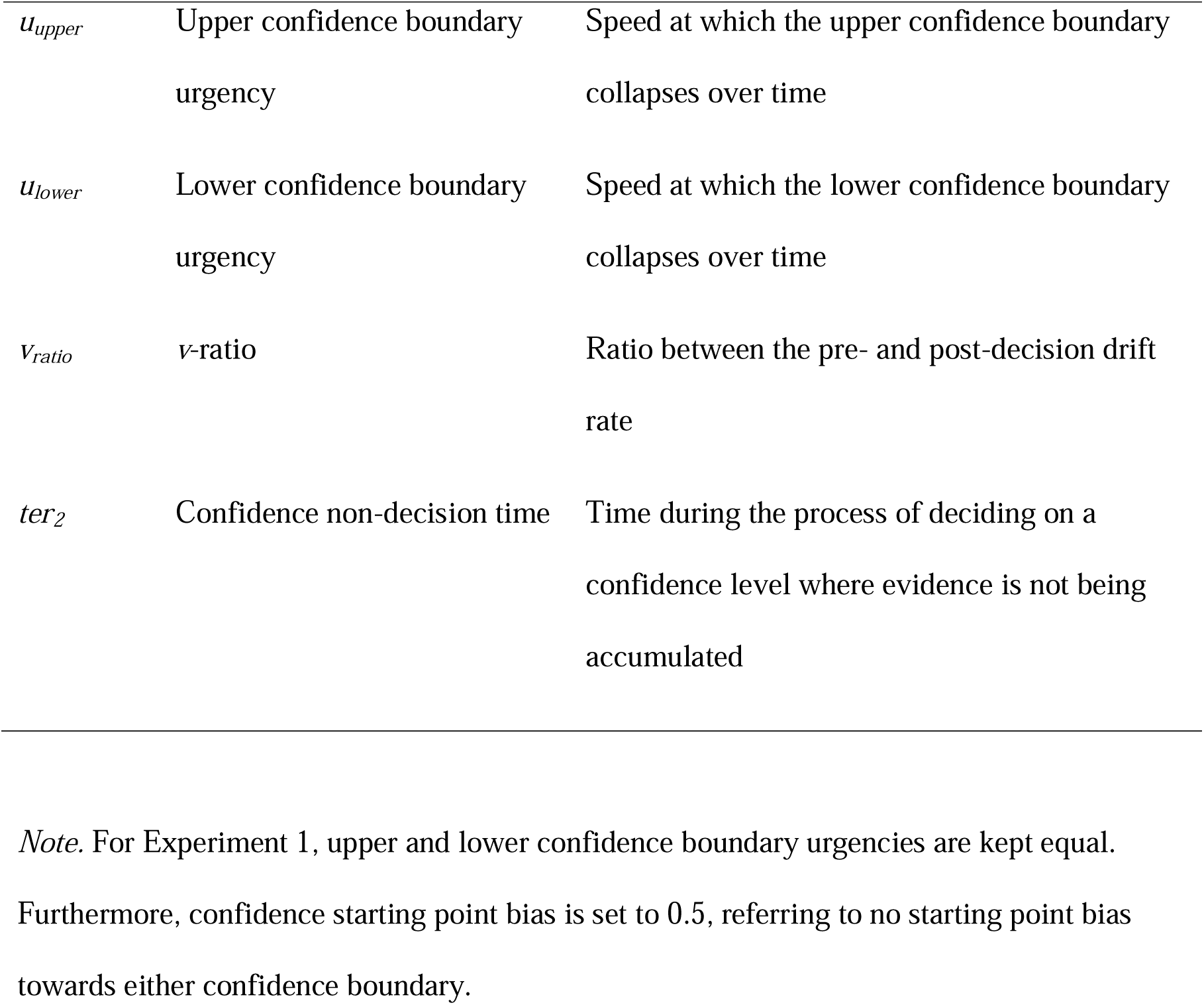
Free parameters of the FCB model.

**Table S4.** Free parameters of the Van Zandt Race model.

| Parameter | Meaning | Description |
| --- | --- | --- |
| $a$ | Decision boundary height | Evidence difference between the starting point and decision boundary |
| $v_{correct}$ | Correct choice drift rate | Average increase of evidence supporting the correct choice |
| $v_{error}$ | Error choice drift rate | Average increase of evidence supporting the error choice |
| $ter$ | Non-decision time | Time during the decision-making process where |
|  |  | evidence is not being accumulated |
| $a_2$ | Confidence boundary height | Evidence difference between the starting point and confidence boundary |
| $ter_2$ | Confidence non-decision time | Time during the process of deciding on a confidence level where evidence is not being accumulated |

**Table S5.**
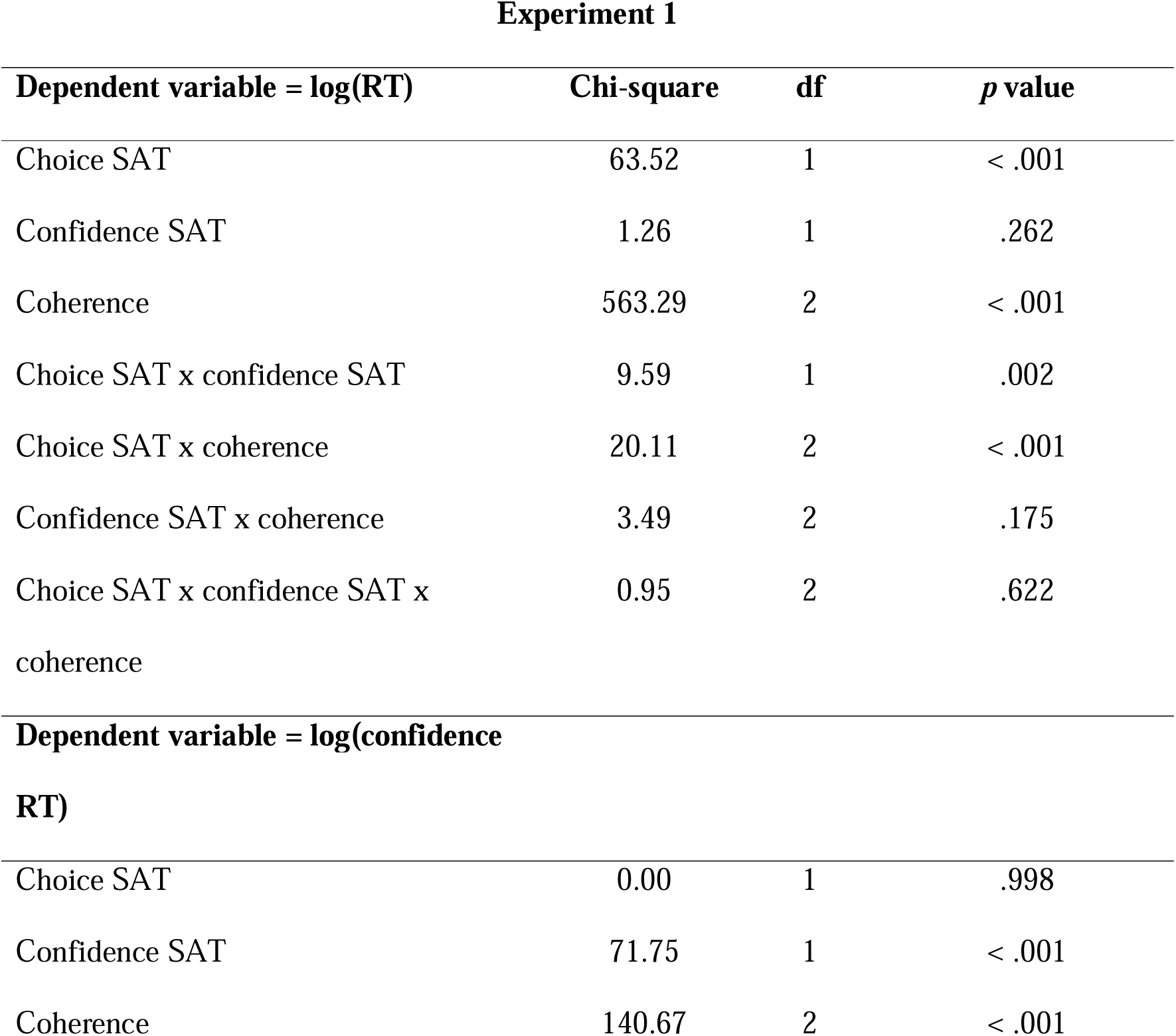
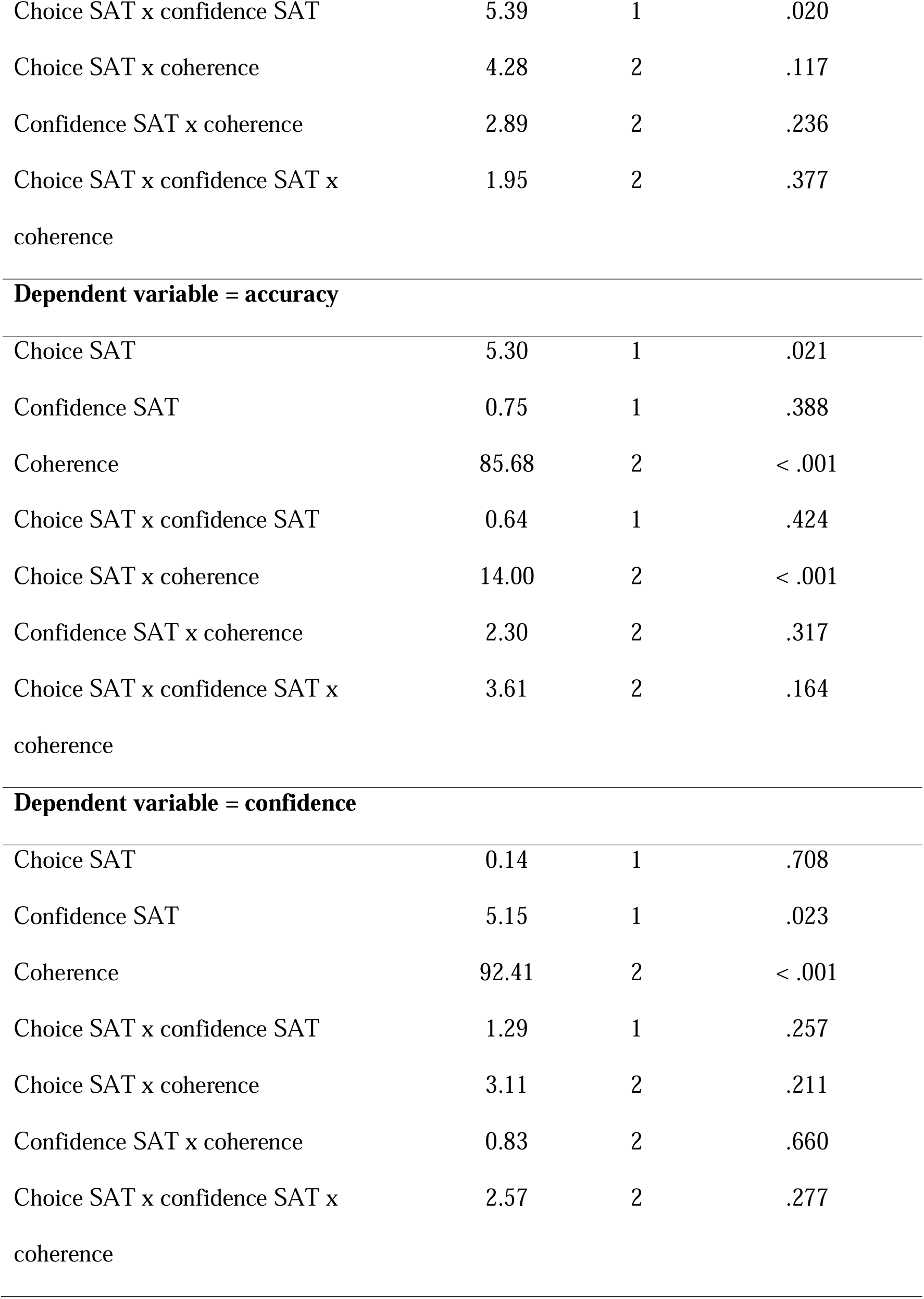
Full results table of the analyses on the FCB model predictions for Experiment 1.

Experiment 1
| Dependent variable = log(RT) | Chi-square | df | *p* value |
| --- | --- | --- | --- |
| Choice SAT | 63.52 | 1 | < .001 |
| Confidence SAT | 1.26 | 1 | .262 |
| Coherence | 563.29 | 2 | < .001 |
| Choice SAT x confidence SAT | 9.59 | 1 | .002 |
| Choice SAT x coherence | 20.11 | 2 | < .001 |
| Confidence SAT x coherence | 3.49 | 2 | .175 |
| Choice SAT x confidence SAT x coherence | 0.95 | 2 | .622 |
| Dependent variable = log(confidence RT) |  |  |  |
| Choice SAT | 0.00 | 1 | .998 |
| Confidence SAT | 71.75 | 1 | < .001 |
| Coherence | 140.67 | 2 | < .001 |
| Choice SAT x confidence SAT | 5.39 | 1 | .020 |
| Choice SAT x coherence | 4.28 | 2 | .117 |
| Confidence SAT x coherence | 2.89 | 2 | .236 |
| Choice SAT x confidence SAT x<br>coherence | 1.95 | 2 | .377 |
| **Dependent variable = accuracy** |  |  |  |
| Choice SAT | 5.30 | 1 | .021 |
| Confidence SAT | 0.75 | 1 | .388 |
| Coherence | 85.68 | 2 | < .001 |
| Choice SAT x confidence SAT | 0.64 | 1 | .424 |
| Choice SAT x coherence | 14.00 | 2 | < .001 |
| Confidence SAT x coherence | 2.30 | 2 | .317 |
| Choice SAT x confidence SAT x<br>coherence | 3.61 | 2 | .164 |
| **Dependent variable = confidence** |  |  |  |
| Choice SAT | 0.14 | 1 | .708 |
| Confidence SAT | 5.15 | 1 | .023 |
| Coherence | 92.41 | 2 | < .001 |
| Choice SAT x confidence SAT | 1.29 | 1 | .257 |
| Choice SAT x coherence | 3.11 | 2 | .211 |
| Confidence SAT x coherence | 0.83 | 2 | .660 |
| Choice SAT x confidence SAT x<br>coherence | 2.57 | 2 | .277 |

**Table S6.**
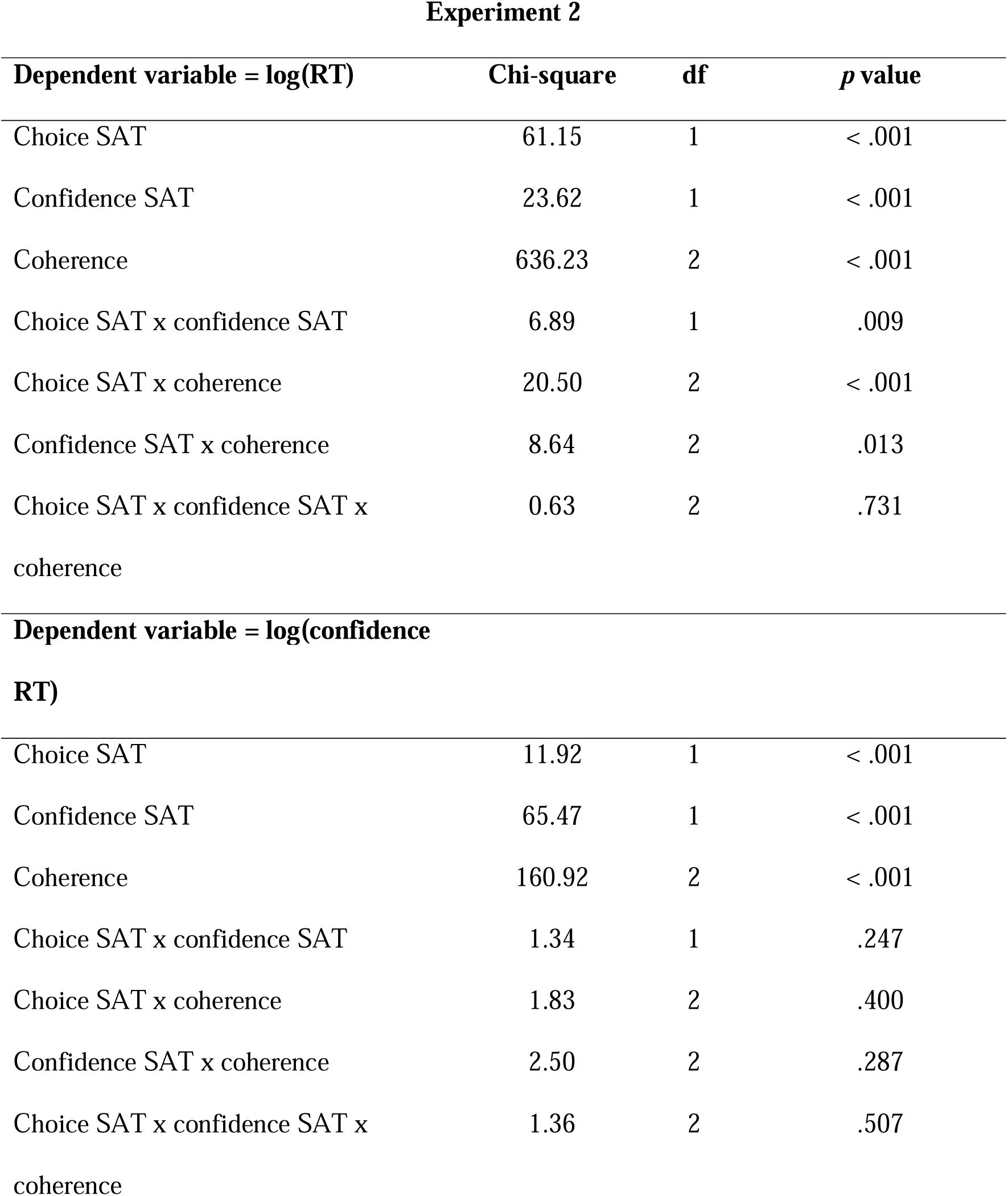
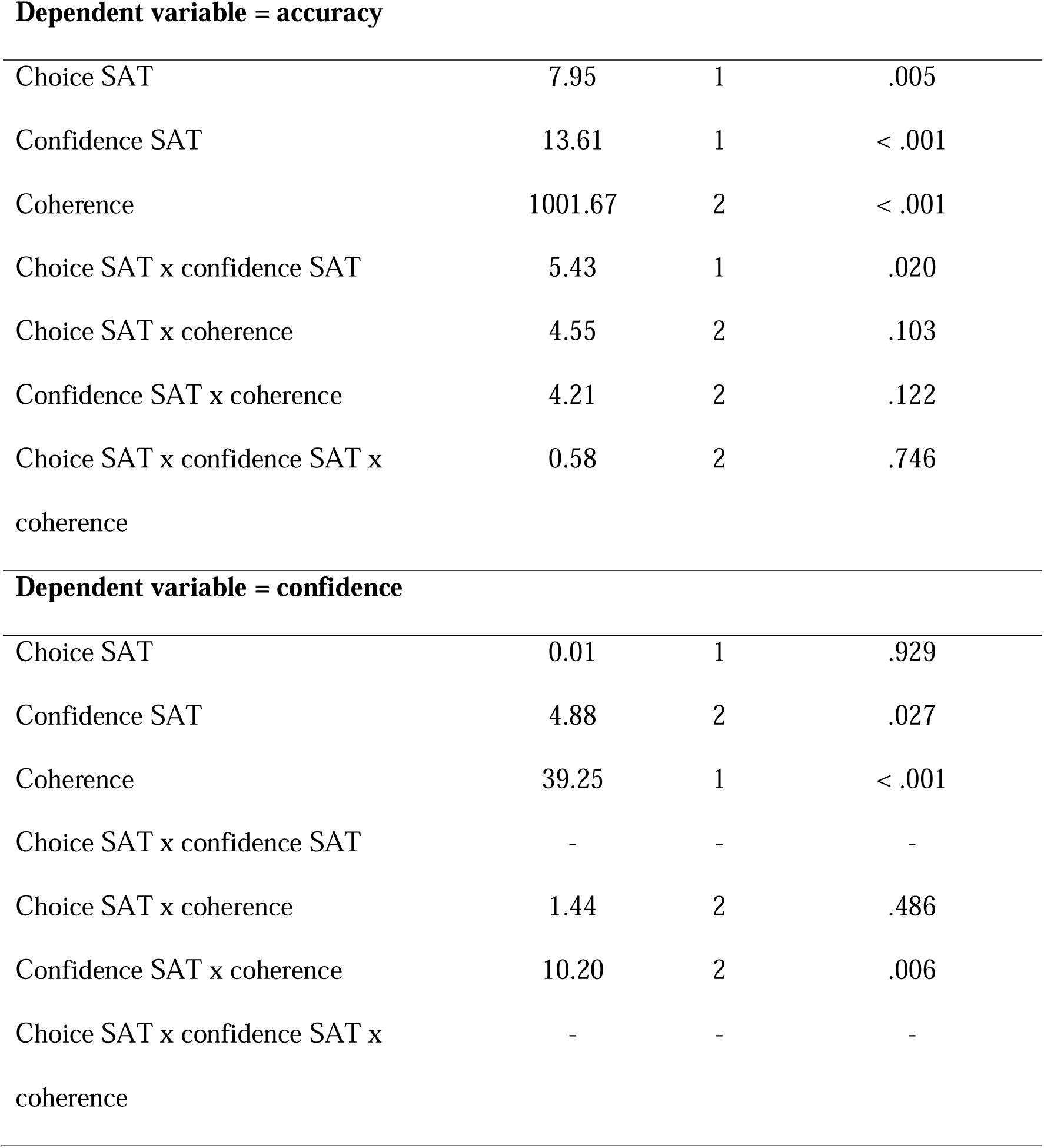
Full results table of the analyses on the FCB model predictions for Experiment 2. Note that two interaction effects were omitted from the confidence model in the behavioral data (due to inflated VIF values), and these were likewise omitted here in order to be able to directly compare both fits.

Experiment 2
| Dependent variable = log(RT) | Chi-square | df | *p* value |
| --- | --- | --- | --- |
| Choice SAT | 61.15 | 1 | < .001 |
| Confidence SAT | 23.62 | 1 | < .001 |
| Coherence | 636.23 | 2 | < .001 |
| Choice SAT x confidence SAT | 6.89 | 1 | .009 |
| Choice SAT x coherence | 20.50 | 2 | < .001 |
| Confidence SAT x coherence | 8.64 | 2 | .013 |
| Choice SAT x confidence SAT x<br>coherence | 0.63 | 2 | .731 |
| **Dependent variable = log(confidence**<br>**RT)** |  |  |  |
| Choice SAT | 11.92 | 1 | < .001 |
| Confidence SAT | 65.47 | 1 | < .001 |
| Coherence | 160.92 | 2 | < .001 |
| Choice SAT x confidence SAT | 1.34 | 1 | .247 |
| Choice SAT x coherence | 1.83 | 2 | .400 |
| Confidence SAT x coherence | 2.50 | 2 | .287 |
| Choice SAT x confidence SAT x<br>coherence | 1.36 | 2 | .507 |
| **Dependent variable = accuracy** |  |  |  |
| Choice SAT | 7.95 | 1 | .005 |
| Confidence SAT | 13.61 | 1 | < .001 |
| Coherence | 1001.67 | 2 | < .001 |
| Choice SAT x confidence SAT | 5.43 | 1 | .020 |
| Choice SAT x coherence | 4.55 | 2 | .103 |
| Confidence SAT x coherence | 4.21 | 2 | .122 |
| Choice SAT x confidence SAT x<br>coherence | 0.58 | 2 | .746 |
| **Dependent variable = confidence** |  |  |  |
| Choice SAT | 0.01 | 1 | .929 |
| Confidence SAT | 4.88 | 2 | .027 |
| Coherence | 39.25 | 1 | < .001 |
| Choice SAT x confidence SAT | - | - | - |
| Choice SAT x coherence | 1.44 | 2 | .486 |
| Confidence SAT x coherence | 10.20 | 2 | .006 |
| Choice SAT x confidence SAT x<br>coherence | - | - | - |

**Figure S1.**
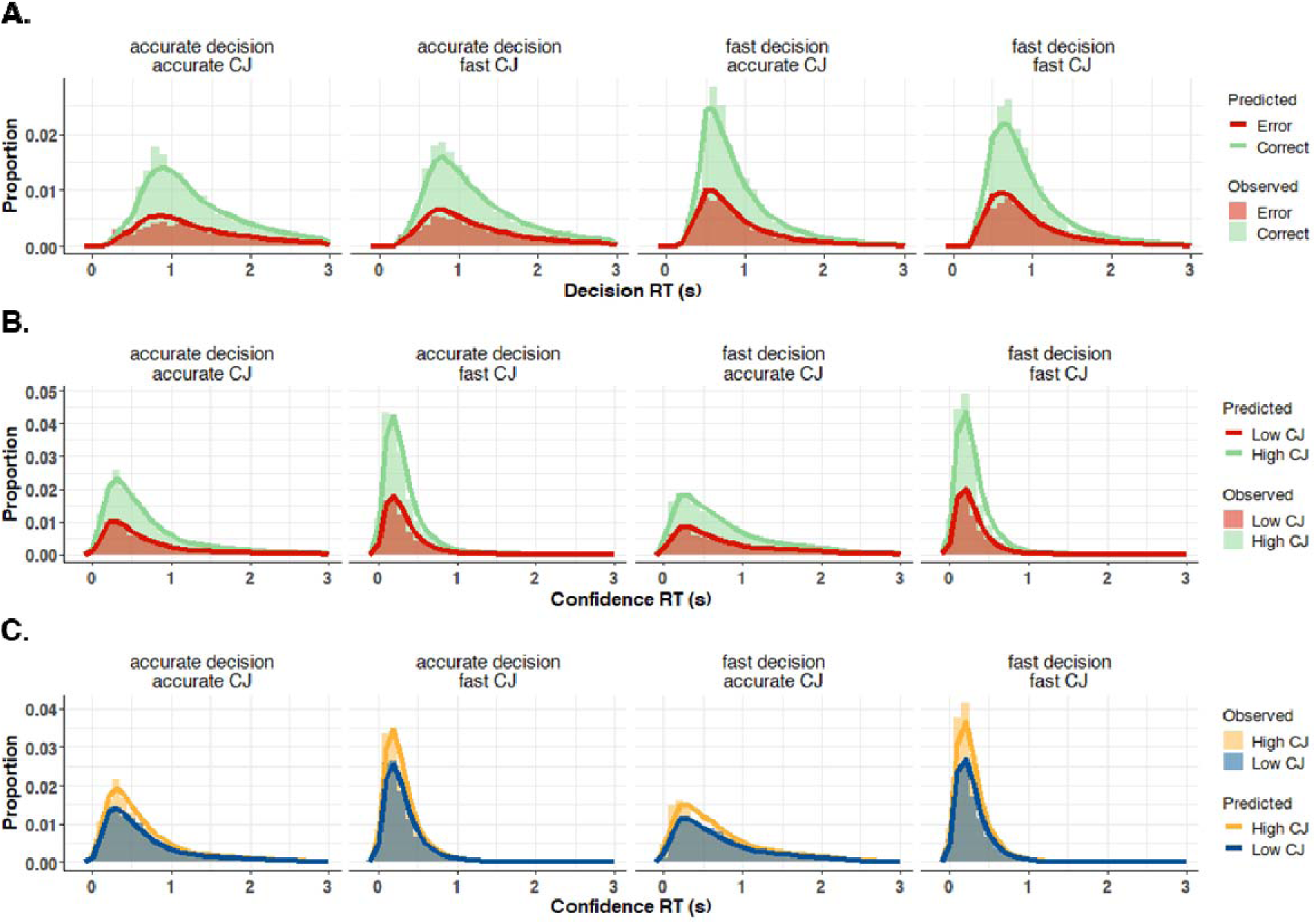
Experiment 1 Model Fit of the FCB model

**Figure S2.**
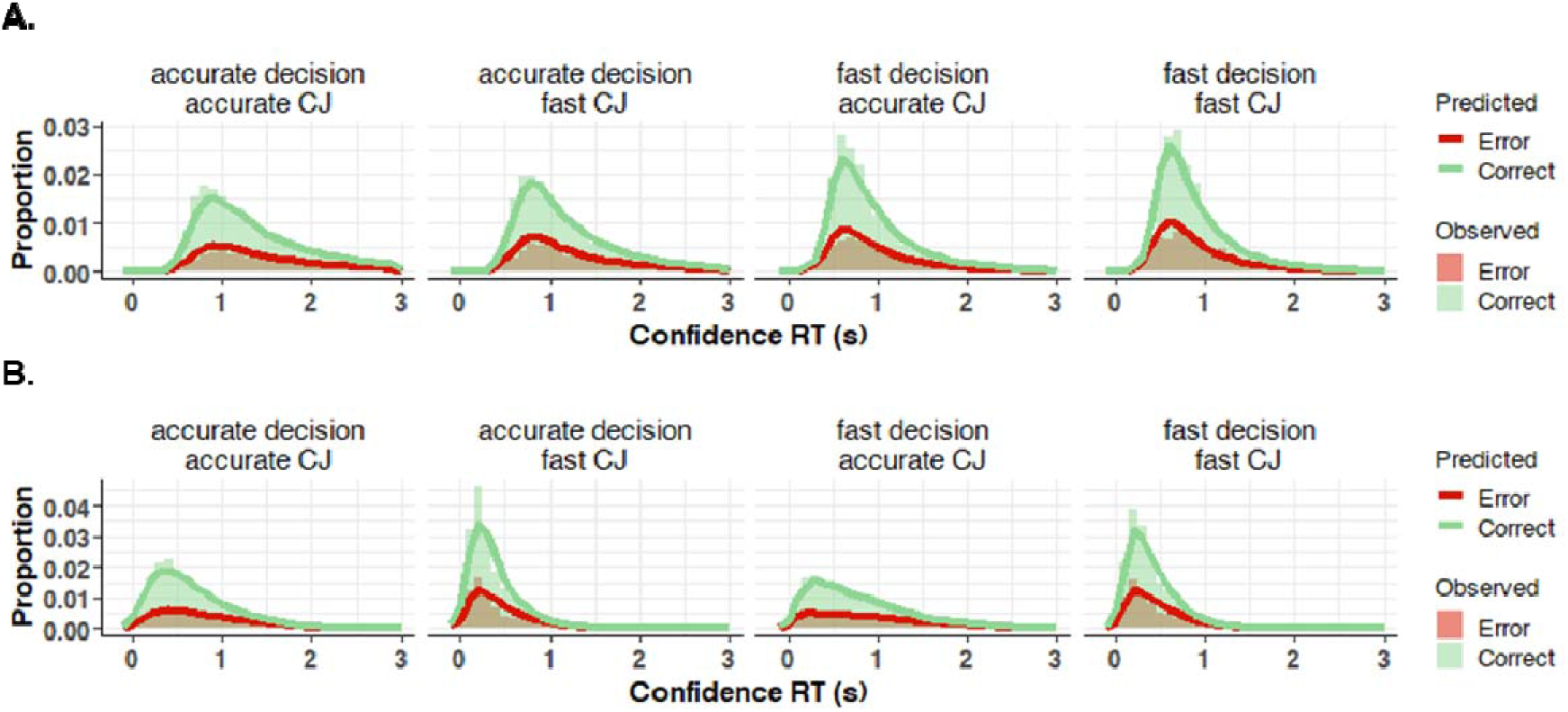
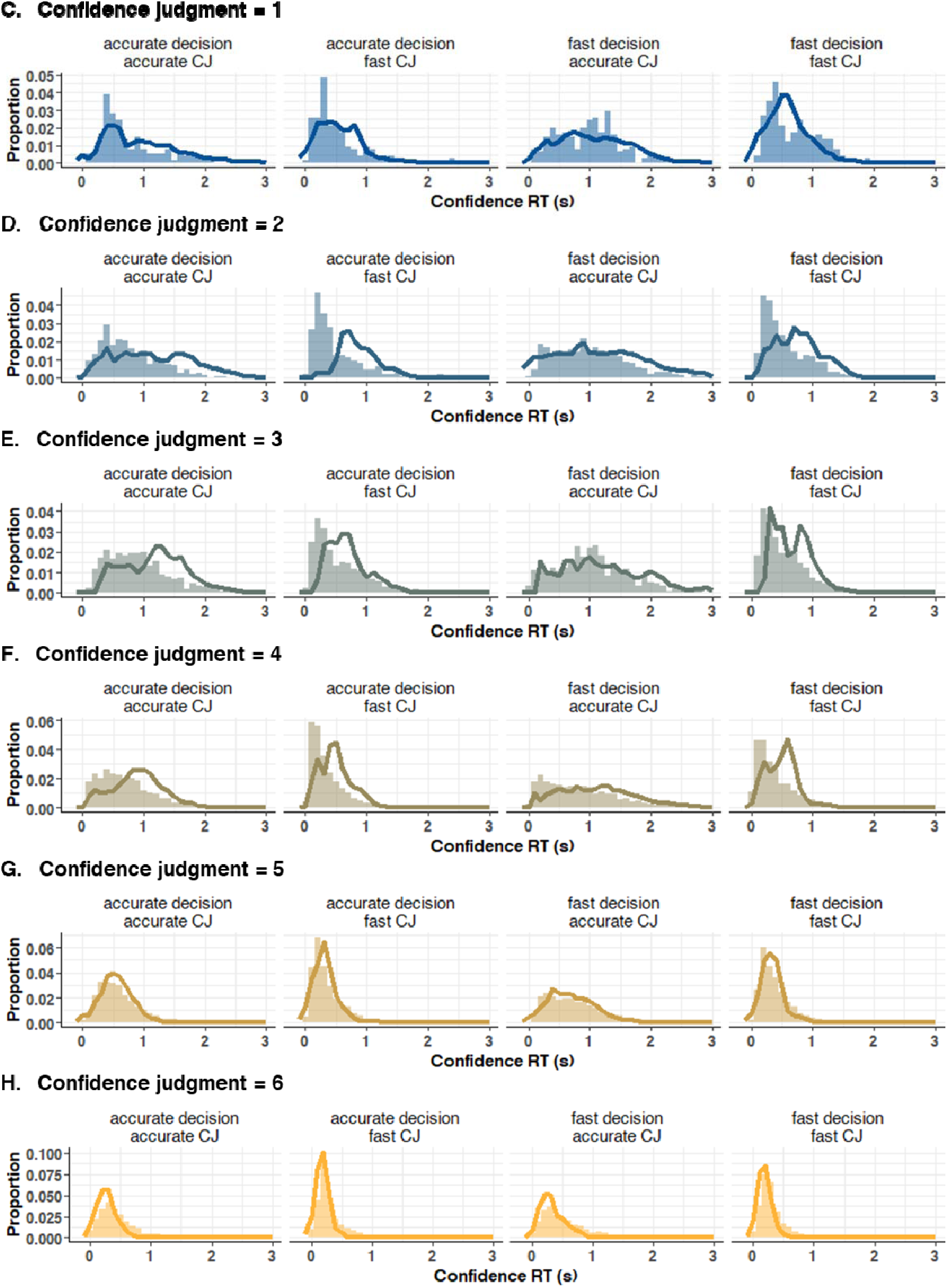
Experiment 2 Model Fit of the FCB model

### Parameter Recovery

We performed a parameter recovery on the best fitting models of Experiment 1 and 2 to ensure the interpretability of the model parameters. To this end, we simulated data for *N* = 100 synthetic participants, once with N_trials_ = 30.000 (i.e., simulating an ideal scenario) and once with N_trials_ = 180 (i.e. the number of trials per cell in our design). Parameters were randomly sampled from a uniform distribution with the minimum and maximum values chosen such that they were in line with the empirically observed fits; For Experiment 1, *a* = [1, 3], *a_2_* = [0, 4], *u* = [0, 1], *v_1,2,3_* = [0, 2], *ter* = [0, 1], *ter_2_*= [-.5, .5], *v_ratio_* = [0, 2]. For Experiment 2, *a* = [1, 3], *a_2_* = [2, 7], *u_upper_* = [0, 3], *u_lower_* = [0, 1.5], *v_1,2,3_* = [0, 2], *ter* = [0, 1], *ter_2_* = [-.5, .5], *v_ratio_* = [0, 1], *z_2_* = [-.5, .5]. Subsequently, we computed the correlation between observed and estimated parameters.

Inspection of the correlation results of Experiment 1 revealed that with 30.000 trials all parameters recovered well (*r_s_*> 0.70), with exception of the urgency parameter, *u,* (*r* = 0.16). As shown in Figure S1, a few urgency parameters did not recover and *u* should therefore be interpreted with caution. This is likely caused by the limited sensitivity of the cost function of Experiment 1 to confidence RT – CJ relationships (i.e. given that there were only two confidence options). Results did not change drastically when the parameter recovery was repeated with 180 trials (Figure S2).

Parameter recovery of Experiment 2 with 30.000 trials showed excellent recovery for all parameters (*r_s_* > 0.91; Figure S3). In line with these results, parameters recovered well with 180 trials (*r_s_* > 0.70; Figure S4). Notably, the inclusion of 6 levels of confidence allowed the model to also recover the urgency parameters.

**Figure S3.**
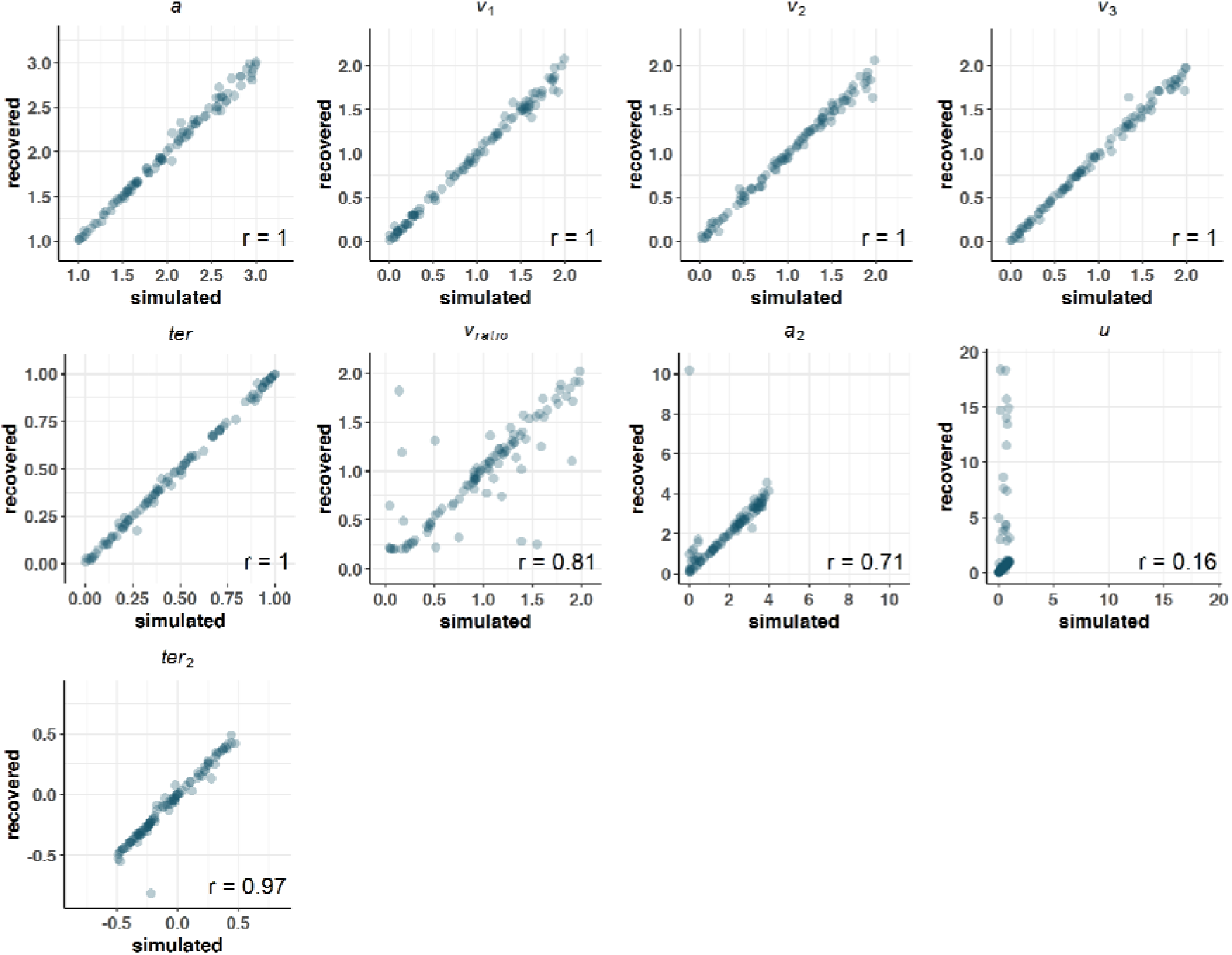
Experiment 1 FCB Parameter Recovery (N_trials_ = 30.000)

**Figure S4.**
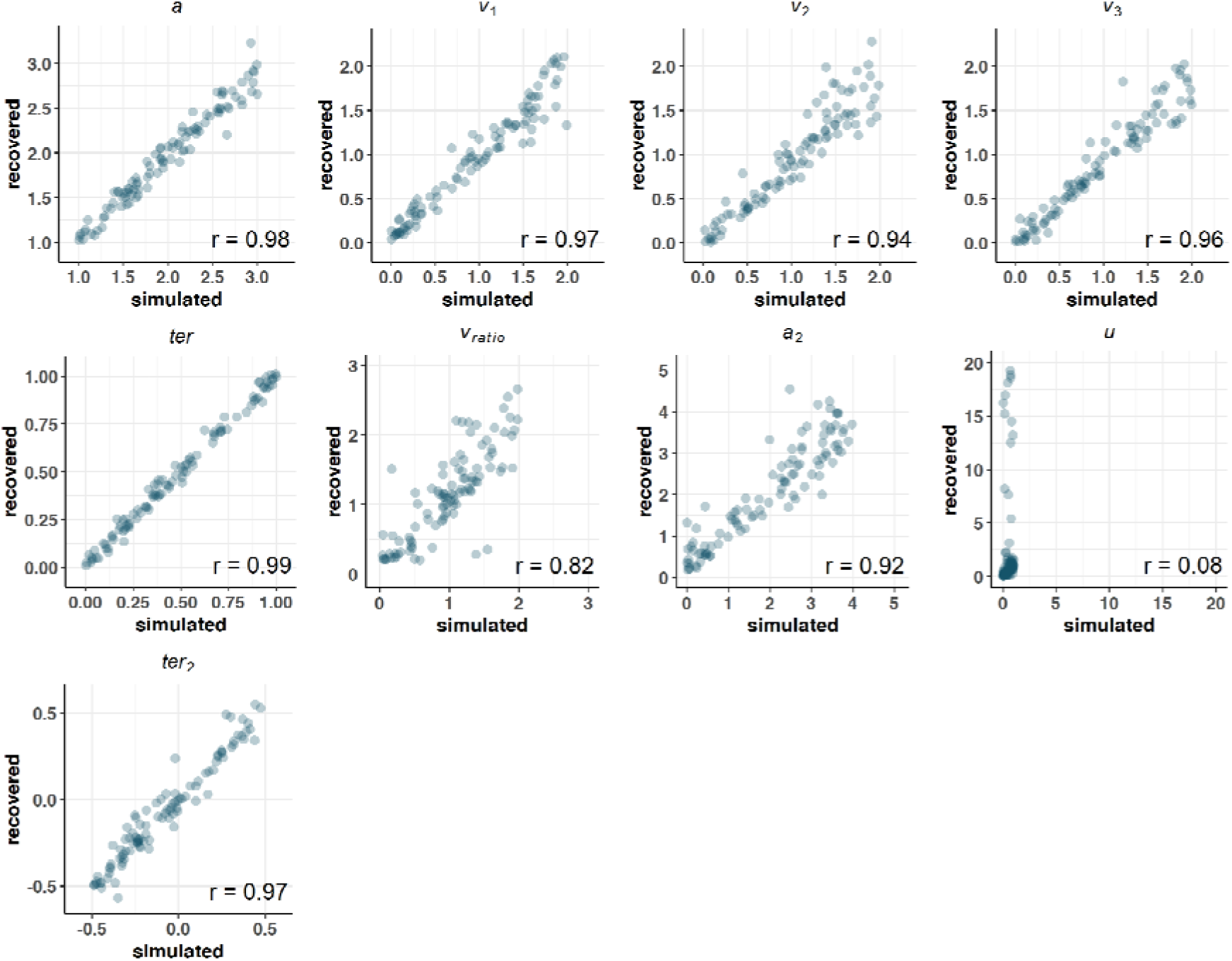
Experiment 1 FCB Parameter Recovery (N_trials_ = 180)

**Figure S5.**
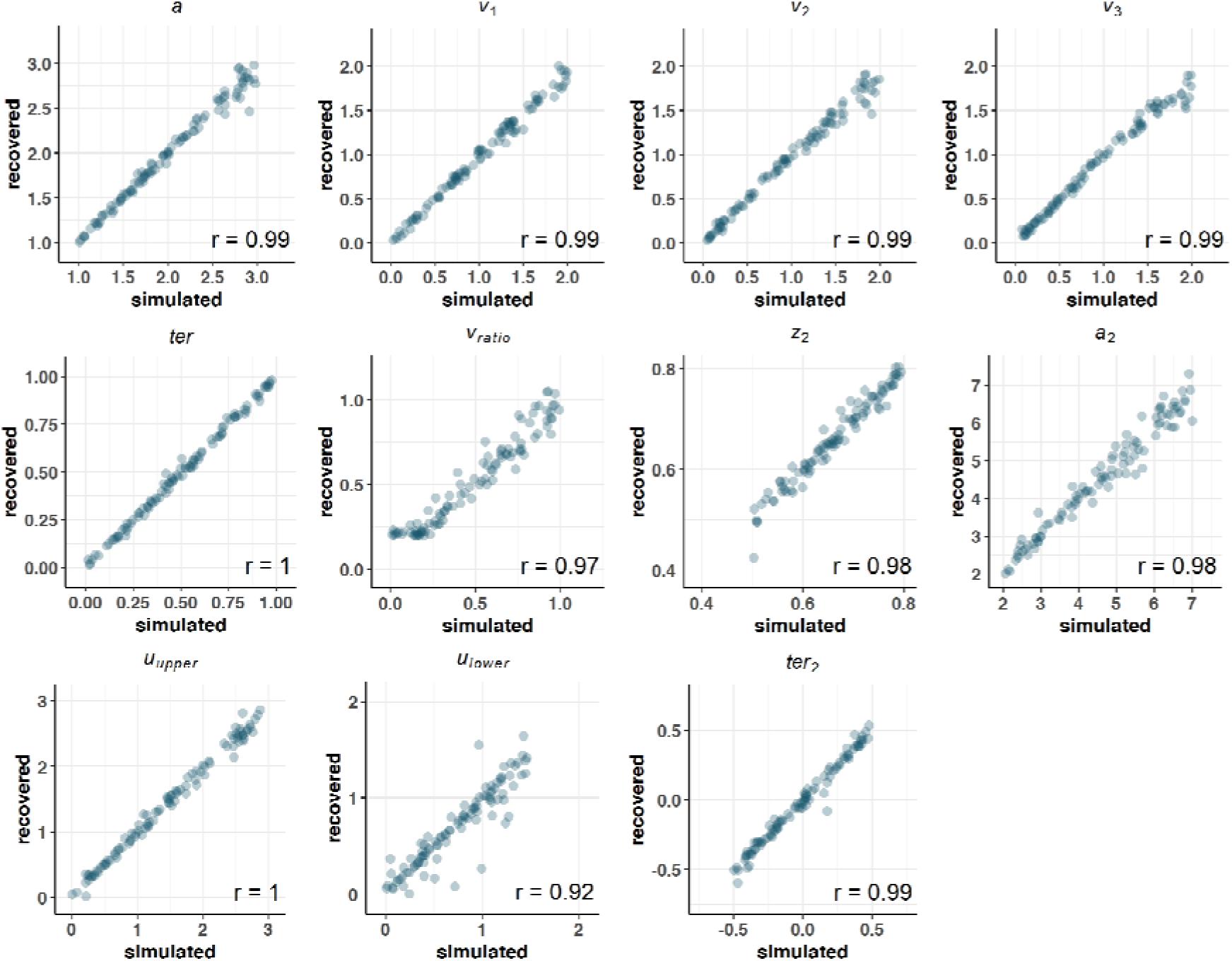
Experiment 2 FCB Parameter Recovery (N_trials_ = 30.000)

**Figure S6.**
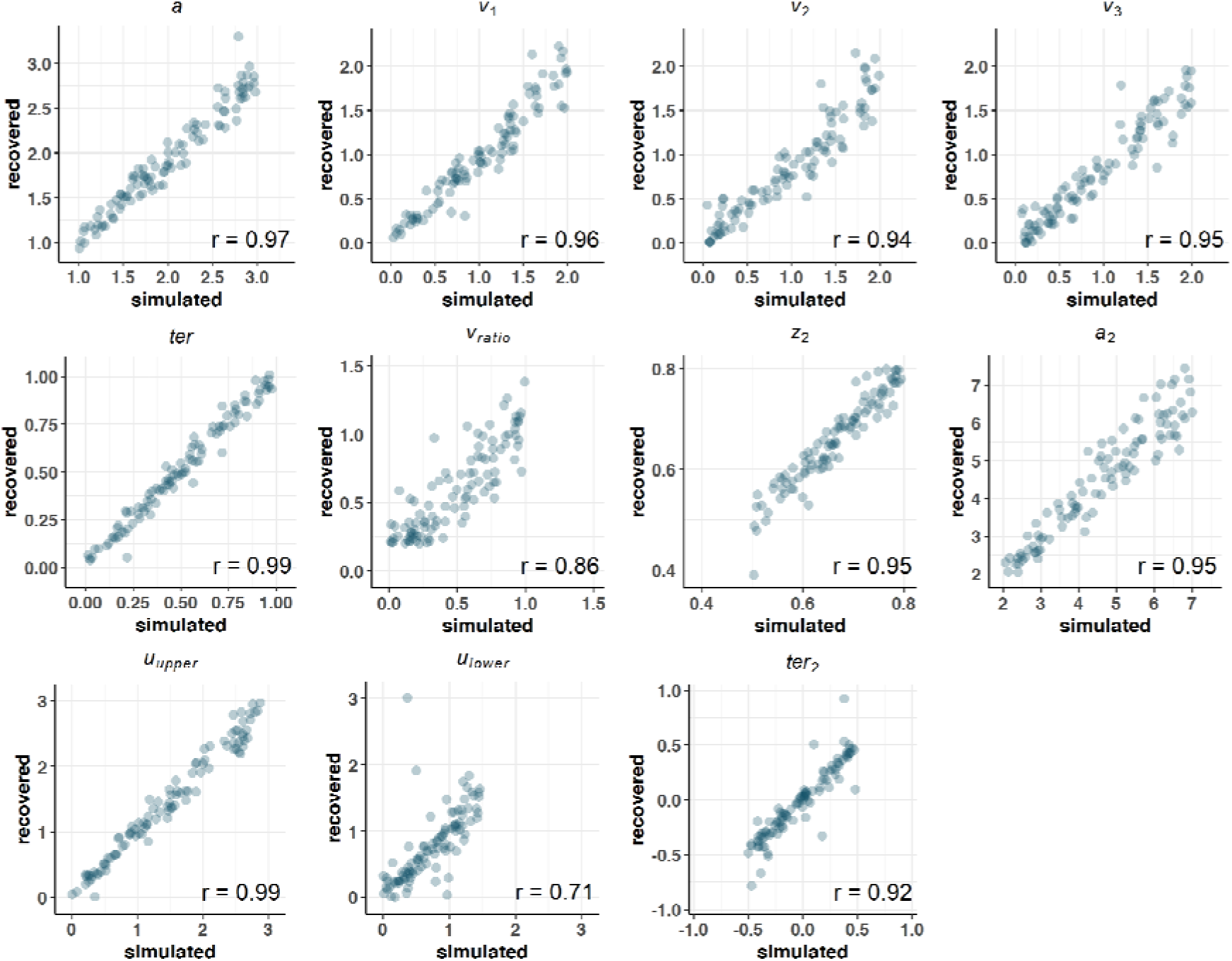
Experiment 2 FCB Parameter Recovery (N_trials_ = 180)

## Notes

https://github.com/StefHerregods/ConfidenceBounds

### Competing Interest Statement

The authors have declared no competing interest.

### Summary of Updates

We have now compared our model to 3 competing models, and showed that it provides the best fit both in quantitative and qualitative terms.

